# Optimising neural contrast and minimising tissue shrinkage in dragon lizard brains: buffered Lugol’s iodine versus phosphotungstic acid for contrast-enhanced µCT

**DOI:** 10.64898/2026.10.13.758998

**Authors:** F. Navea, J.R. Black, S. Whiteley, D.S.B. Dissanayake, C.C. Kerr, S.P. Collin, J.M. Crowe-Riddell

## Abstract

Contrast-enhanced micro-computed tomography (µCT) has become a valuable tool for addressing neuroanatomical questions that previously required destructive histology. As a relatively new technique, it remains subject to ongoing technical refinement, and taxon- and tissue-specific protocol optimisation. We evaluated the performance of two common contrast-enhancing agents, buffered Lugol’s iodine and phosphotungstic acid (PTA) in aqueous solution, for µCT visualisation of the nervous system in the Jacky dragon lizard, *Amphibolurus muricatus* by comparing staining time, contrast specificity and brain tissue shrinkage evaluated at three concentrations (1.25%, 2.5% and 5%). Buffered Lugol’s iodine outperformed aqueous PTA by requiring shorter staining times (as short as four days) and providing better white matter-grey matter differentiation, whereas PTA required longer incubation periods of approximately three weeks and produced poorer white-grey matter differentiation. Importantly, buffered Lugol’s iodine produced higher brain-to-endocast occupancy estimates than aqueous PTA, suggesting substantially reduced tissue shrinkage. PTA-treated specimens exhibited an estimated 13.3% reduction in occupancy and approximately 20% greater brain tissue shrinkage than buffered Lugol’s-treated specimens. Together, these findings demonstrate that buffered Lugol’s iodine staining provides the high level of internal tissue contrast required to resolve fine-scale neuroanatomical organisation in three dimensions, preserves the spatial relationships among anatomical structures, and reduces artefacts, all of which result in a closer correspondence between the brain and endocranial cavity. From a practical perspective, the study provides guidance for contrast agent selection in future µCT studies and reveals that buffered Lugol’s iodine is better suited to fine-scale neuroanatomical, volumetric and geometric morphometric analyses, whereas PTA is advantageous for applications prioritising general soft-tissue or musculoskeletal visualisation.

**Graphical Abstract:** Buffered Lugol’s iodine improves µCT visualisation of dragon lizard brains through faster staining, greater neural contrast, reduced tissue shrinkage and closer correspondence between brain and endocast surfaces compared with phosphotungstic acid (PTA).

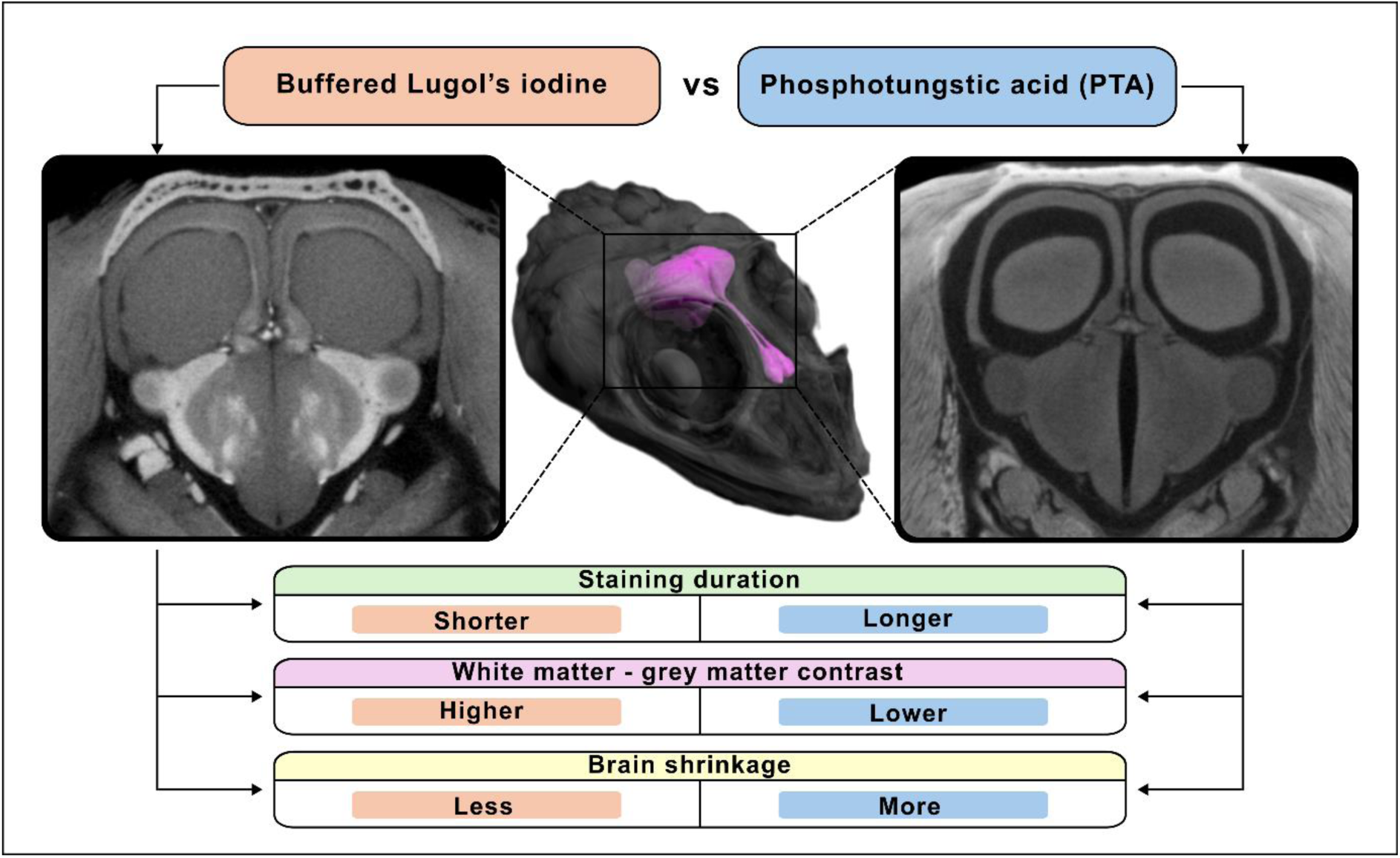

## Introduction

Our understanding of the anatomy of biological systems, including the nervous system, advances alongside technological and methodological developments in bioimaging. As imaging tools become more precise and accessible, technical innovations improve our ability to characterise biological structures accurately and to address novel research questions. In recent decades, non-invasive, three-dimensional (3D) bioimaging techniques, such as magnetic resonance imaging (MRI), X-ray computed tomography (CT), light sheet fluorescence microscopy (LSFM), and optical projection tomography (OPT), have significantly advanced neuroanatomical and morphological research. By enabling the visualisation of anatomical structures across multiple levels of biological organisation, these approaches have improved the quantification of cellular components, the description of spatial relationships among brain structures, and the characterisation of neural connectivity in diverse taxa, both *in vivo* and *ex vivo* (Dawood *et al*., 2021; Gignac and Kley, 2018).

Compared with classical histological methods, which require destructive and time-consuming procedures such as dissection, embedding, sectioning, and staining, three-dimensional bioimaging offers advantages for non-destructive *in situ* characterisation of soft tissues. Although histology generally provides high cellular resolution, tissue processing can introduce morphological artefacts and compromise the preservation of three-dimensional spatial relationships. Consequently, bioimaging approaches are particularly valuable for broad comparative studies and for the examination of rare or irreplaceable museum specimens, including specimens of endangered species that cannot be destructively sampled (Collin *et al*., 2026).

X-ray computed tomography, and particularly micro-computed tomography (µCT), has emerged as a complementary and often more accessible alternative to MRI for high-resolution anatomical imaging. µCT is a rapid and relatively inexpensive technique that uses X-rays projected at multiple angles around the specimen. It enables the differentiation of tissues based on the degree of radiation absorbed by the sample, which is detected and quantified by the scanner (Bushberg *et al*., 2012; Lauridsen *et al*., 2011). While µCT provides high-resolution imaging for mineralised and calcified tissues such as bones and cartilage, it does not naturally yield strong contrast in soft tissues, such as the brain and nervous system, that absorb X-rays poorly (Mizutani and Suzuki, 2012; Neues and Epple, 2008). This limitation can be overcome by treating samples with contrast agents containing elements with high atomic numbers that penetrate and accumulate in biological tissues, thereby enhancing X-ray absorption and generating contrast comparable to that of dense structures like bone or cartilage. With enhanced contrast staining, µCT produces non-destructive, high-resolution and high-contrast images of animal soft tissues (Gignac *et al*., 2016; Metscher, 2009a). Moreover, staining procedures are relatively simple, as most contrast agents penetrate the neural tissues via passive diffusion in solution, contributing to the rapid increase in studies employing this technique. For a recent comprehensive review of contrast agents used in µCT, see Balcaen *et al*. (2025).

As a relatively new technique, contrast-enhanced µCT remains subject to ongoing methodological refinement and protocol optimisation. Despite being simple and effective, it has several limitations: most contrast agents induce osmotic or chemical tissue shrinkage; stain penetration can be slow or uneven in large specimens and staining may be irreversible. Furthermore, image quality is highly sensitive to fixation and storage conditions, stain type and concentration, the pH and osmolarity of the solution, and the size of the specimen.

For high-resolution neuroanatomical imaging using contrast-enhanced µCT, two contrast agents have become the most widely used for studying the brains of vertebrates and invertebrates across both embryonic and adult stages: inorganic iodine (I₂) and phosphotungstic acid (hereafter referred to as PTA). Both agents are suitable for staining freshly formaldehyde-fixed and ethanol-preserved specimens, as they can be dissolved in aqueous or ethanolic solutions. This versatility likely contributes to their frequent use in recent museum-based studies, expanding data availability across a broad range of taxa and developmental stages, including both cartilaginous and bony fishes (Camilieri-Asch *et al*., 2020a; Clement *et al*., 2015; Deary *et al*., 2016; Kolmann *et al*., 2023; Staggl *et al*., 2023), amphibians (Krings *et al*., 2017; Naumann and Olsson, 2018), reptiles (Callahan *et al*., 2021; Collin *et al*., 2026; Hughes *et al*., 2016; Zhou *et al*., 2025), birds (Straight *et al*., 2024; Watanabe *et al*., 2019), mammals (Carlisle *et al*., 2017; Gignac *et al*., 2021; Mrzílková *et al*., 2019) and invertebrates (Rivera-Quiroz and Miller, 2022; Rother *et al*., 2021; Sakurai and Ikeda, 2019; Smith *et al*., 2016). However, the selection of a contrast agent is often driven by limited studies on the target species, commercial availability, anecdotal evidence of effectiveness or, in some cases, trial and error. This is particularly relevant because, despite their similar applicability for soft tissue staining, iodine and PTA exhibit distinct tissue affinities attributable to their respective chemical properties, which can lead to important differences in performance and suitability for different applications.

Iodine, commonly used as inorganic iodine (I₂) in alcohol or as Lugol’s iodine (I₂KI) in aqueous solution, shows affinity for lipids and carbohydrates present in soft (neural) tissues, allowing discrimination between myelinated and unmyelinated nervous tissue (Gignac and Kley, 2018). Iodine also diffuses comparatively rapidly and, although it produces discolouration of the sample, this change is potentially reversible (Callahan *et al*., 2021; Gignac *et al*., 2021; Lanzetti and Ekdale, 2021). However, it has the disadvantage of inducing soft tissue shrinkage depending on concentration, an effect that can be mitigated using a recently developed buffered Lugol’s iodine solution protocol (Dawood *et al*., 2021). Owing to its widespread use, iodine-based approaches have been formalised as diffusible iodine-based contrast-enhanced computed tomography (diceCT; Gignac *et al*., 2016). Iodine preferentially binds to lipids creating enhanced contrast of neural tracts that contain myelin (e.g. in vertebrate brains) (Gignac and Kley, 2018). Conversely, PTA, commonly dissolved in ethanol or water, binds preferentially to proteins and connective tissue (Metscher, 2009b; Rivera-Quiroz and Miller, 2022). Although PTA requires longer penetration times, it has been reported to induce less soft tissue shrinkage than non-buffered iodine-based stains when dissolved in water (Buytaert *et al*., 2014) and is widely applied in invertebrate neuroanatomical studies.

In vertebrates, iodine has been shown to improve differentiation between white and grey matter, potentially facilitating the identification of a range of brain features such as nuclei, layers, fibres and tracts, whereas PTA has been reported to better preserve soft-tissue volume, which may be advantageous for morphometric analyses. Comparative assessments of the performance of iodine and PTA as contrast-enhancing agents for µCT imaging of nervous tissue remain limited, and most of these studies have been conducted in alcoholic or non-buffered aqueous solutions, before buffered Lugol’s was shown to mitigate shrinkage in liver tissue by Dawood *et al*. (2021). Consequently, a direct comparison of buffered Lugol’s iodine and PTA is lacking, limiting our understanding of their effects on soft tissues and whether they can be used interchangeably in comparative neuroanatomical studies. Evaluating their staining efficacy and ability to preserve nervous-tissue morphology during imaging is therefore essential for the accurate assessment of the nervous system.

Reptiles remain comparatively understudied in neuroscience relative to other vertebrates such as fishes, birds, and mammals. Their high species diversity and extensive representation in museum collections make them an ideal taxon for comparative neuroanatomical and morphological studies. Here, we evaluate buffered Lugol’s iodine and aqueous PTA as contrast agents for µCT imaging in reptiles using the Jacky dragon, *Amphibolurus muricatus* (Squamata: Agamidae) as a model species. This charismatic species has been well-studied for its social communication and visual acuity (Carlile *et al*., 2006; Peters and Evans, 2003; Peters *et al*., 2007). Jacky dragons are also representative of the ecologically diverse Australian agamid radiation (Powney *et al*., 2010), which is an emerging group for understanding the impacts of temperature and ecology on morphology and brain evolution (Esquerré *et al*., 2014; Harlow and Taylor, 2000; Hoops *et al*., 2017a, 2017b).

## Materials and Methods

### Specimen collection and preparation

A total of 15 specimens of the adult Jacky dragon (*Amphibolurus muricatus)* were sourced from a captive colony at the University of Canberra (n = 4) or collected from Buckley Park Foreshore Reserve and Point Lonsdale, Victoria, Australia (n = 11), between December 2021 and January 2022. The majority of specimens were male (n = 13), and all specimens were of a similar size (snout to vent length: 78–104 mm; mass: 20.82–39.87 g). Animals were euthanised, decapitated and their heads fixed via immersion in 4% paraformaldehyde in 0.1 M phosphate buffer (PFA). All procedures adhered to University of Canberra animal ethics guidelines (Project 270) and complied with national welfare guidelines. Specimens were then shipped to La Trobe University, Bundoora, Victoria, Australia, for brain tissue analyses.

### Experimental design

Specimens were assigned to two contrast-enhanced µCT staining treatments, buffered Lugol’s iodine and PTA, which formed the primary comparison of this study. Each treatment comprised multiple staining conditions varying in concentration and, for PTA, solution pH. Treatment conditions, specimen allocation, and subsequent analyses are summarised in Table 1. A subset of specimens was additionally processed for histological validation. Prior to staining, all specimens were manually skinned to facilitate stain penetration, as the highly cornified reptilian epidermis can impede contrast agent diffusion (Collin *et al*., 2026).

**Table 1.** Specimens and corresponding treatments specifications. C: contrast quantification, H: histology, P: pH-adjusted PTA staining, S: staining time assessment, V: Volumetric and morphological analyses.

| Specimen | Sex | SVL (mm) | Weight (g) | Staining agent | Concentration | pH | Experiment |
| --- | --- | --- | --- | --- | --- | --- | --- |
| AmLT1 | Male | 98 | 36.73 | Buffered Lugol's iodine | 1.25% | 7.2 | S, C, V |
| AmLT7 | Male | 85 | 27.28 | Buffered Lugol's iodine | 1.25% | 7.2 | V |
| AmLT2 | Male | 103 | 39.48 | Buffered Lugol's iodine | 2.50% | 7.2 | S, C, V |
| AmLT3 | Male | 100 | 39.87 | Buffered Lugol's iodine | 5% | 7.2 | S, C, V |
| 9512 | Female | 80 | 22.08 | Aqueous PTA | 1.25% | 2 | S, C, V |
| 81821 | Male | 82 | 24.01 | Aqueous PTA | 1.25% | 2 | V |
| 81928 | Male | 85 | 24.01 | Aqueous PTA | 2.50% | 2 | V |
| AmLT8 | Male | 78 | 20.29 | Aqueous PTA | 2.50% | 2 | S, C, V |
| 81820 | Female | 83 | 21.03 | Aqueous PTA | 5% | 2 | S, C, V |
| AmLT12 | Male | 89 | 28.88 | Aqueous PTA | 1.25% | 7.2 | P |
| AmLT14 | Male | 81 | 23.98 | Aqueous PTA | 1.25% | 7.2 | P |
| AmLT4 | Male | 86 | 20.82 | Aqueous PTA | 1.25% | 4 | P |
| AmLT5 | Male | 104 | 35.07 | Aqueous PTA | 2.50% | 4 | P |
| AmLT6 | Male | 96 | 29.06 | Cresyl violet | 0.0143% | 4 | H |
| AmLT10 | Male | 94 | 35.23 | Cresyl violet | 0.0143% | 4 | H |

#### Buffered Lugol’s iodine staining

Buffered Lugol’s staining (Lugol’s iodine prepared in Sorensen’s buffer) followed established diceCT protocols for soft-tissue imaging (Callahan *et al*., 2021; Gignac *et al*., 2016), with buffering incorporated following Dawood *et al*. (2021) to minimise staining-induced tissue shrinkage. To evaluate the effects of staining concentration on penetration rate, contrast quality, and tissue integrity, specimens were immersed in buffered Lugol’s at one of three concentrations (1.25%, 2.5%, or 5.0% w/v) and incubated at room temperature in the dark on a rotary shaker. These concentrations were selected to represent a commonly used range for Lugol’s iodine staining in diceCT studies (Gignac *et al*., 2016; Vickerton *et al*., 2013). Staining solutions were replaced whenever they became visibly faded or increasingly transparent, indicating depletion of iodine through uptake by the specimen.

#### PTA staining

PTA staining followed established protocols for soft-tissue imaging (De Greef *et al*., 2015; Descamps *et al*., 2014). Skinned head specimens were immersed in aqueous PTA prepared in Type II laboratory-grade purified water at one of three concentrations (1.25%, 2.5%, or 5.0% w/v) and incubated at room temperature in the dark on a rotary shaker. These concentrations were selected to encompass the range commonly used for PTA staining in contrast-enhanced µCT studies (De Greef *et al*., 2015; Descamps *et al*., 2014), while extending the evaluation to a higher concentration (5.0% w/v) to explore its effects on stain penetration, contrast quality, and tissue integrity. PTA solutions were prepared in purified water without buffering to follow established (previously published) staining protocols to permit direct evaluation of this widely used methodology and were replaced weekly. Additional specimens were included to provide an exploratory qualitative assessment of the influence of solution pH on staining performance. For this purpose, 1.25% and 2.5% PTA solutions were adjusted to selected pH values using NaOH (Table 1).

#### Histology

Selected specimens were processed for standard histology to provide neuroanatomical reference material. Following fixation, brains were carefully dissected from the brain cavity, embedded in 14% (w/v) gelatin from porcine skin Type A (Merck/Sigma-Aldrich, St. Louis, MO, USA), sectioned at 40 µm using a cryostat (Leica CM1850 cryostat; Leica Microsystems, Wetzlar, Germany), and stained with 0.0143% (w/v) cresyl violet acetate in aqueous acetate buffer (pH ≈ 4.0). Sections were mounted, coverslipped using DPX, and digitised using a slide scanner (Zeiss

Axioscan 7 slide scanner; Carl Zeiss Microscopy GmbH, Jena, Germany) using a 10x objective. Histological sections were used to validate the anatomical correspondence and tissue differentiation observed in the contrast-enhanced µCT datasets, particularly the distinction between white and grey matter.

### Micro computed tomography scanning and image analysis

Prior to scanning, all specimens were rinsed in phosphate-buffered saline (PBS 0.01 M) for 1 h, wrapped in gauze and placed in 20 mL vials filled with PBS to prevent specimen movement and dehydration during image acquisition. Scans were conducted at the Trace Analysis for Chemical, Earth and Environmental Sciences (TrACEES) facility at The University of Melbourne. µCT scanning was performed with a Phoenix Nanotom m scanner (Waygate Technologies, Wunstorf, Germany) operated using xscontrol and Phoenix Datos|x acquisition software (Waygate Technologies, Wunstorf, Germany). Scanning parameters are listed in Supplementary File S1. Volume reconstruction of the µCT data was performed using Phoenix datos|x reconstruction software (Waygate Technologies, Wunstorf, Germany) by applying an inline median filter, an ROI filter and a beam hardening correction filter during reconstruction. Reconstructed volumes were exported as TIFF image stacks and imported into Dragonfly 3D World Software version 2025.1 (Comet Technologies Canada Inc., Montreal, QC, Canada) for segmentation, 3D visualisation and assessment of staining performance. For each specimen, an initial scan was performed without staining to obtain images without soft-tissue contrast, providing a bone reference for easier endocast segmentation. Subsequently, multiple periodic scans were conducted to document staining progress and contrast intensity. Staining time, image contrast, and tissue shrinkage were evaluated across all concentration-series.

### Stain penetration assessment

Stain penetration time was quantified using one specimen from each staining concentration for each staining agent (n = 3 per staining agent) as the number of days required for the staining agent to reach the innermost regions of the brain. Penetration was assessed by visually inspecting the progression of the staining front from the specimen surface towards its centre across successive scans. All specimens were evaluated using the same coronal virtual section at the level of the posterior telencephalon/anterior diencephalon. This plane of section was selected because it contains the largest cross-sectional area of brain tissue, providing a representative view of stain progression. For consistency, the coronal plane was defined as perpendicular to the rostro-caudal axis of the head. Complete stain penetration was defined as the first scan in which no background-intensity voxels remained within the innermost regions of the brain, indicating that the staining agent had reached the entire brain volume. This was further confirmed by inspecting the entire brain volume in addition to the selected coronal section.

### Tissue contrast quantification

Using the same specimens, greyscale values of the background, bone, white matter, grey matter, endocranial space, and muscle were measured at five time points for buffered Lugol’s-stained specimens and six time points for PTA-stained specimens throughout the staining period. Reconstructed TIFF stack datasets from each specimen were manually aligned in Dragonfly, and the Point set tool within the Annotate toolbox was used to place six probe points at predefined anatomical locations on a coronal virtual slice at the level of the optic chiasm (Supplementary Fig. S1). This level allowed all tissue classes of interest to be sampled within a single section. Probe point greyscale values were extracted from all aligned datasets, ensuring that identical three-dimensional locations were sampled throughout the staining time series (Supplementary File S2).

To quantitatively evaluate staining performance, tissue contrast was assessed across tissue interfaces relevant to brain visualisation and manual segmentation. These included internal neural differentiation between white matter (WM) and grey matter (GM), separation of the brain from the skull, delineation of the brain from the endocranial cavity, and discrimination between neural and adjacent extracranial soft tissues. Accordingly, Michelson contrast coefficients were calculated between (1) white matter and grey matter, (2) bone and the higher-intensity neural tissue (WM or GM), (3) the lower-intensity neural tissue and the endocranial space, and (4) the higher-intensity neural tissue and muscle (Supplementary File S2). The higher- and lower-intensity neural tissues were determined from the sampled greyscale values at each scanning time point. To enable comparisons among specimens, staining concentrations, and scanning time points, contrast was calculated using the Michelson contrast coefficient:

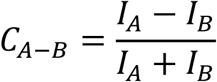

where *C_A_*_−*B*_ is the contrast coefficient between two tissues, and *I_A_* and *I_B_* are their respective greyscale values. For simplicity, these four Michelson contrast coefficients are hereafter referred to as C1: WM minus GM (internal neural contrast), C2: Bone minus higher intensity neural tissue (bone-brain contrast), C3: lower-intensity neural tissue minus endocranial space (brain–endocranial space contrast), and C4: higher-intensity neural tissue minus muscle (brain–muscle contrast). Positive and negative values indicate the direction of the intensity difference, whereas the absolute magnitude indicates the degree of contrast.

An optimal internal neural contrast was defined as the scanning time at which C1 reached its maximum following stain penetration of both sampling locations. Early coefficient peaks resulting from one of the sampled locations remaining unstained were interpreted as artefacts of differential stain diffusion rather than true tissue differentiation and were therefore not considered representative of optimal image contrast.

### Brain occupancy, tissue shrinkage and brain-endocast correspondence

Volumetric analyses were performed on four buffered Lugol’s-treated specimens and five PTA-treated specimens representing the selected concentration series. Brain, brain tissue, and endocast volumes were quantified to evaluate staining-induced changes in brain occupancy of the endocranial cavity and to estimate the degree of brain neural tissue shrinkage. The brain, as the Central Nervous System organ, comprised all intracranial brain structures, including the ventricular system, hypophysis, and pineal and parapineal organs, but excluding the meninges. Brain tissue, as an approximation to the neural parenchyma, comprised the segmented brain volume after exclusion of the ventricular system. The endocast represented the volume enclosed by the endocranial cavity, with its boundaries defined by the inner surface of the skull or, where bone was absent, by the dura mater (Allemand *et al*., 2023; Balanoff *et al*., 2016). The posterior limit of the endocast was defined at the level of the foramen magnum, by drawing a transverse plane between the most posterior margins of the supraoccipital and basioccipital bones in the midsagittal plane.

Brain, ventricular system, and endocast were manually segmented in Dragonfly for each specimen, and their volumes (in mm³) were calculated from voxel counts and known voxel dimensions. Brain tissue volume was calculated by subtracting ventricular volume from total brain volume. Brain-to-endocast ratio (BER), brain tissue-to-endocast ratio (BTER) and total ventricular percentage were calculated as:

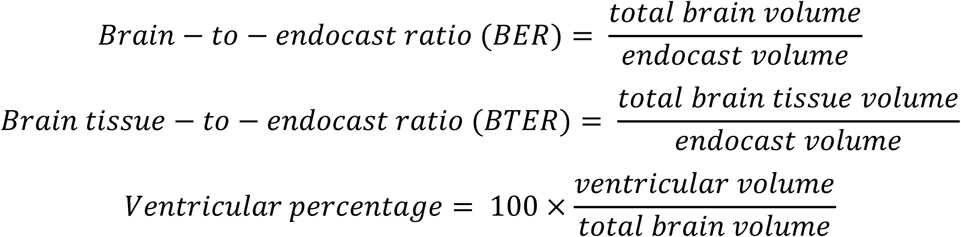

BER quantifies the proportion of the endocranial cavity occupied by the brain, whereas BTER quantifies the proportion occupied by brain tissue. Together, these ratios provide complementary, relative measures of brain occupancy within the endocranial cavity, distinguishing shrinkage of the brain as an organ from shrinkage of the brain tissue independent of ventricular volume. To assess potential effects of specimen size, differences between contrast agents were evaluated using ordinary least squares linear regression models, adjusting separately for body mass, snout–vent length (SVL), head width, or endocast volume (Supplementary File S3).

To further investigate the differences in brain occupancy of the endocranial cavity between staining methods, brain and endocast surface meshes were converted into point clouds and compared in CloudCompare (version 2.14.beta) using a cloud-to-cloud (C2C) distance analysis. In addition to the global volumetric measures provided by BER and BTER, the C2C analysis enabled spatial assessment of brain–endocast correspondence across the entire brain and endocast surfaces. The brain point cloud served as the reference, and absolute nearest-neighbour distances between the brain and endocast point clouds were calculated. Distances were visualised as colour-coded heat maps using a blue-to-red colour scale, with blue indicating close brain–endocast correspondence and red indicating greater separation. The frequency distribution of absolute nearest-neighbour distances was also exported for quantitative comparison between treatments.

## Results and Discussion

Buffered Lugol’s iodine and PTA both enabled high-resolution visualisation of the brain in *Amphibolurus muricatus,* confirming that contrast-enhanced µCT can resolve vertebrate neuroanatomical organisation with a high degree of structural fidelity (Fig. 1). However, the two staining agents differed markedly in their overall performance. Across the concentration range evaluated (1.25%, 2.5%, and 5.0%), buffered Lugol’s iodine consistently produced superior white-grey matter differentiation and less soft-tissue shrinkage than PTA.

**Figure 1.**
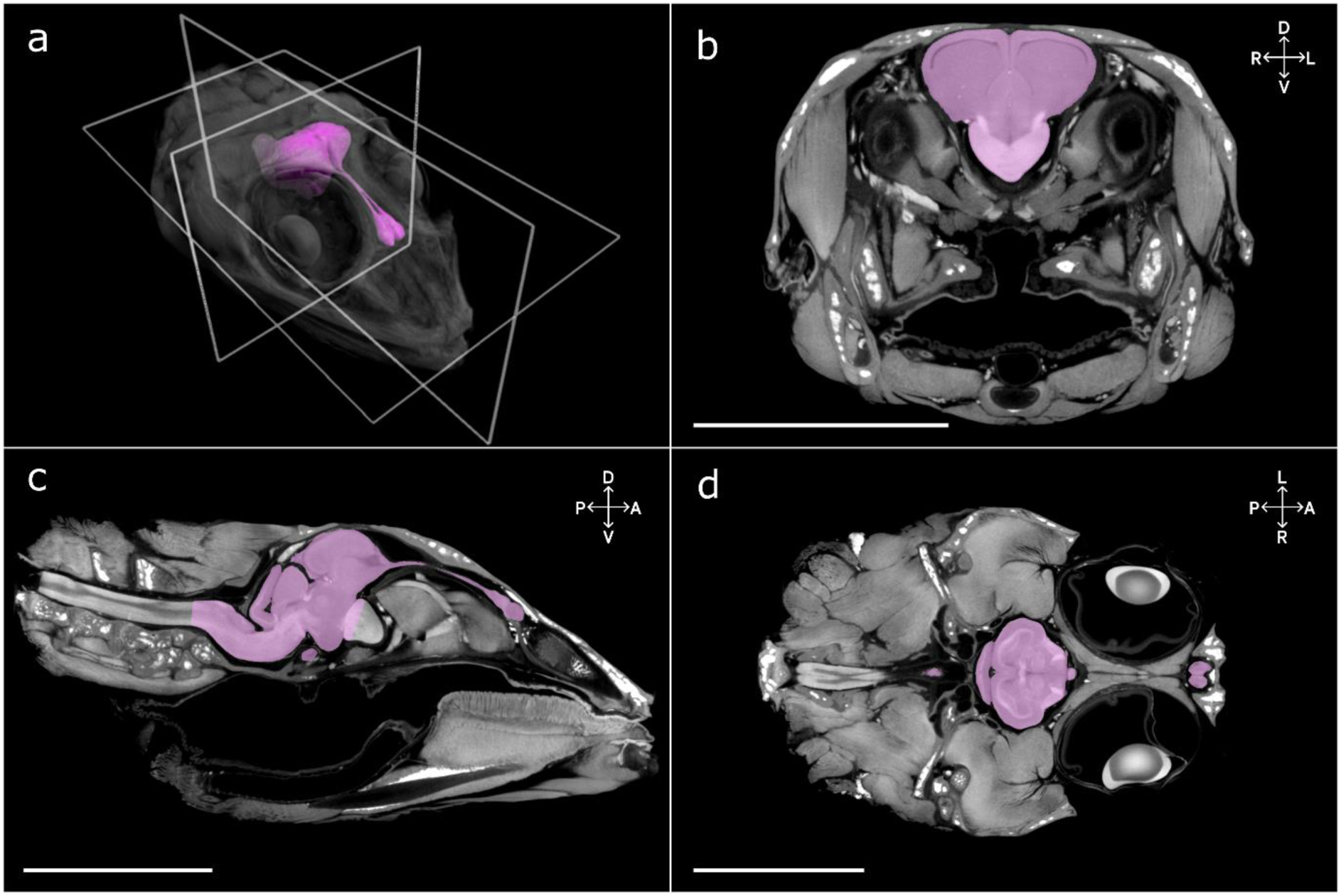
A virtual segmentation of the brain of *A. muricatus*, including a three-dimensional brain model and visualisations in the three principal anatomical planes: b, coronal; c, sagittal, d, horizontal. Soft tissue was enhanced using 5% buffered Lugol’s iodine staining. Scale bars = 1 cm.

### Stain penetration assessment

Stain penetration varied across buffered Lugol’s iodine concentrations, with the lower concentration (1.25%) requiring longer to fully permeate the tissue (Fig. 2). At this concentration, stain diffusion progressed gradually from the peripheral tissues towards deeper structures, with the brain first showing detectable staining after 4 days and complete penetration being observed after 13 days. At higher concentrations, staining progressed more rapidly. In the 2.5% treatment, the brain was almost fully stained after 4 days, with complete penetration occurring between 4 and 8 days. At 5%, complete staining was observed after 4 days. In the specimens examined, increasing concentration above 2.5% produced little additional reduction in the observed time to complete penetration.

**Figure 2.**
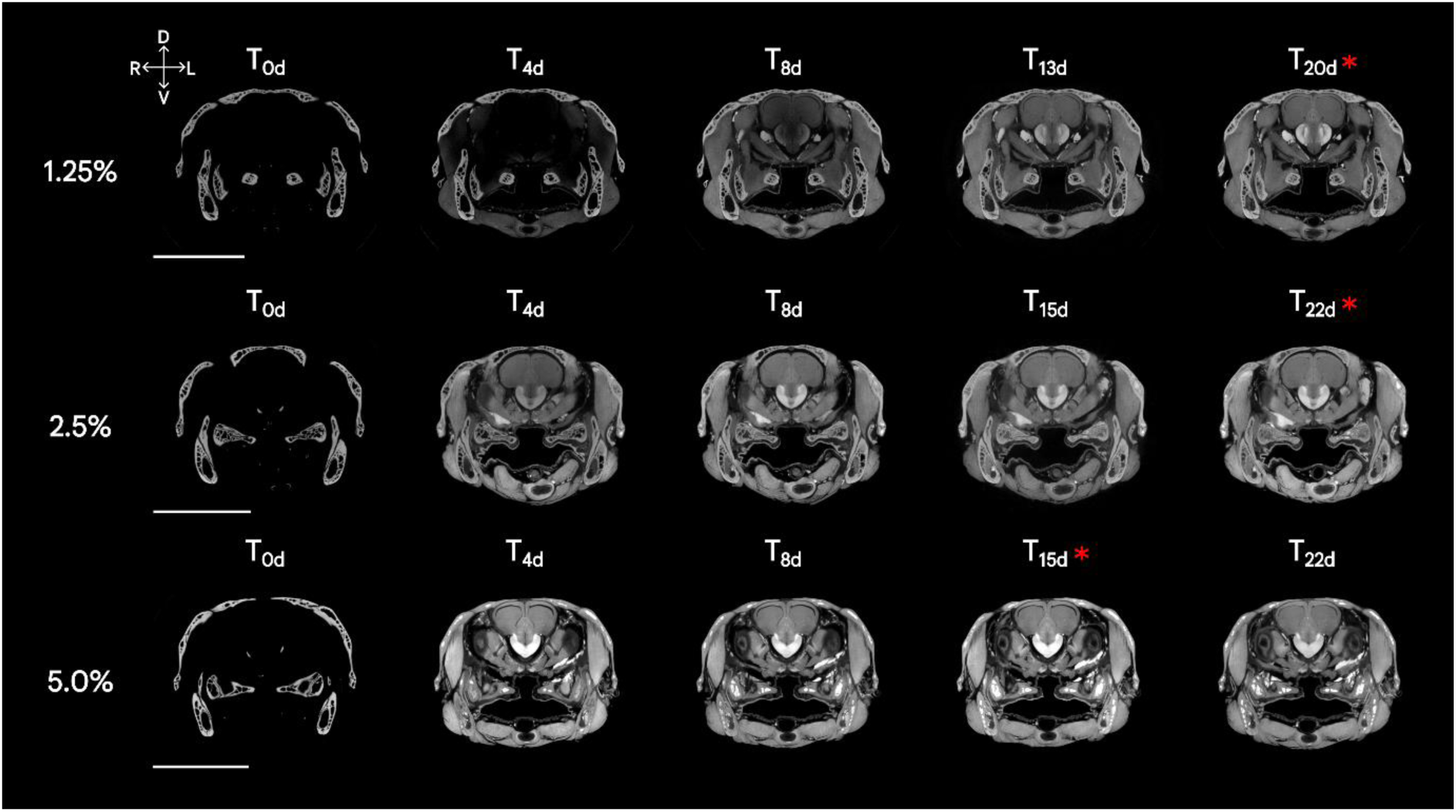
Representative virtual coronal sections of *A. muricatus* heads showing the progression of buffered Lugol’s iodine staining over time at three concentrations (1.25%, 2.5% and 5%). Higher concentrations achieved complete staining quicker, although no differences in peak neural tissue contrast were found between treatments. Time points (T) indicate days post staining. Scanning times at which contrast between white matter and grey matter reached its maximum are marked with an asterisk. Scale bars =1 cm.

PTA exhibited a similar concentration-dependent pattern during the initial stages of staining but progressed considerably more slowly through the specimen overall (Fig. 3). The highest concentration (5.0%) showed the fastest initial uptake, with the staining front advancing more rapidly through the superficial tissues during the first week than in the lower concentrations. Nevertheless, this early difference did not translate into faster complete penetration. Instead, all PTA concentrations required at least 21 days to fully stain the brain, indicating that concentration primarily affected the initial rate of diffusion rather than the total staining time.

**Figure 3.**
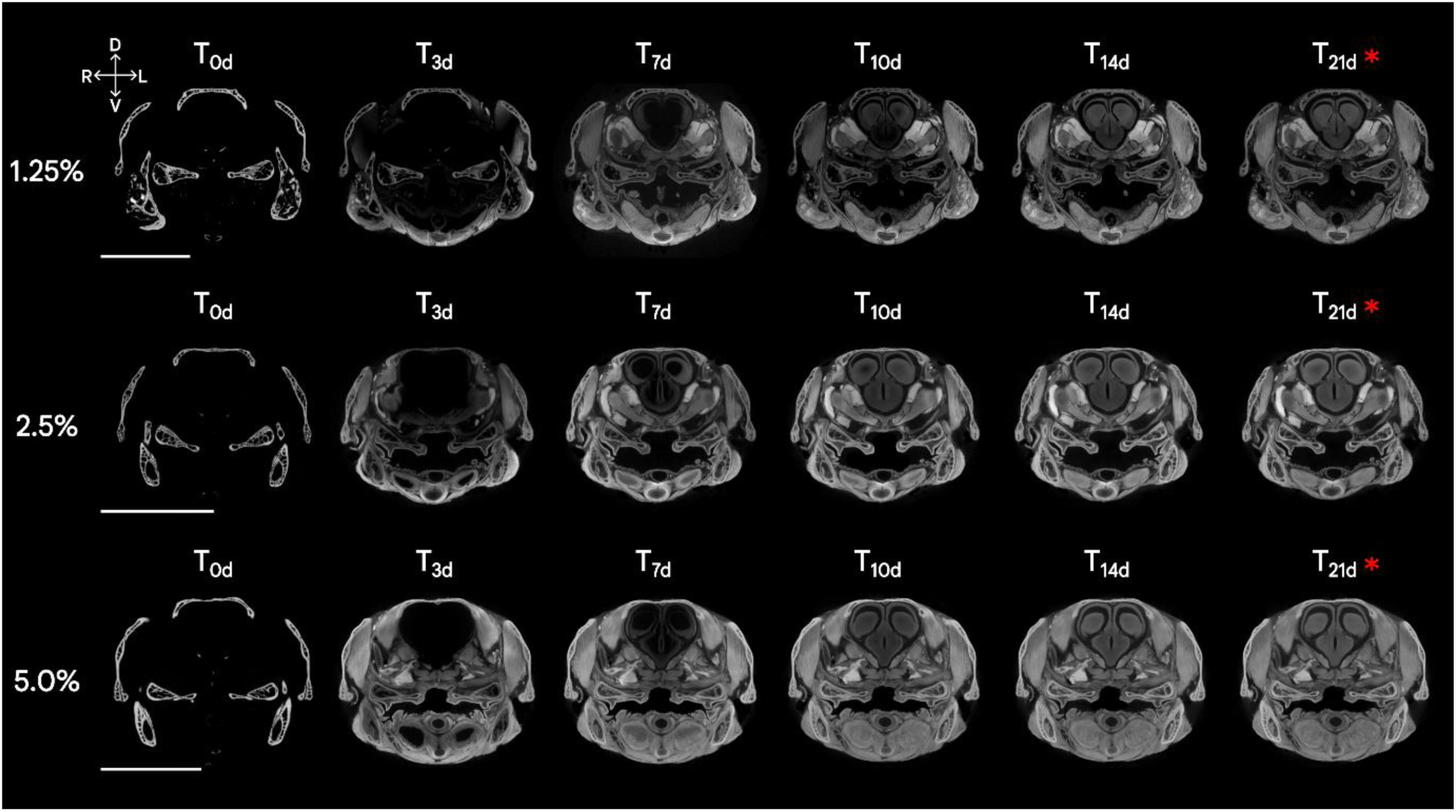
Representative virtual coronal sections of *A. muricatus* heads showing time-dependent PTA staining at three concentrations (1.25%, 2.5%, 5%). Overall soft tissue contrast increases with staining duration, improving brain delineation from the endocranial space, while differentiation between white and grey matter remains limited. Scanning times at which contrast between white matter and grey matter reached its maximum are marked with an asterisk. Scale bars=10mm.

The comparatively longer staining times observed for PTA are consistent with previous studies reporting slower diffusion than iodine-based staining solutions, such as Lugol’s iodine or iodine in ethanol (Chen *et al*., 2018; Descamps *et al*., 2014; Kwon *et al*., 2022; Lesciotto *et al*., 2020; Pauwels *et al*., 2013; Procházková *et al*., 2026; Sakurai and Ikeda, 2019). This difference has generally been attributed to the higher molecular weight of PTA, which limits its diffusion through soft tissues. While our results support the conclusion that PTA requires longer staining times than iodine-based stains, the absolute staining times reported here were likely shortened by our specimen preparation. The use of skinned, isolated heads removed a major barrier to stain entry and provided direct posterior access to the cranial cavity following decapitation, likely facilitating stain penetration. Consequently, the staining times reported here may underestimate those expected for intact specimens, while more accurately reflecting preparation protocols for studies focusing on the central nervous system.

Additionally, buffered Lugol’s iodine solutions were replaced once or twice per week based on visual assessment of solution colour, which provided a practical indicator of stain depletion. As PTA solutions are essentially colourless, an equivalent visual criterion could not be applied, resulting in solutions replaced less frequently (weekly) and constituting an additional factor that may have influenced the staining times observed.

### Tissue contrast quantification

The two contrast agents differed most markedly in their ability to resolve internal brain architecture, particularly the distinction between white and grey matter, as well as the contrast between the brain and surrounding musculature, likely reflecting the distinct tissue affinities of iodine and PTA. These differences became more pronounced as staining progressed.

Optimal neural staining duration for buffered Lugol’s iodine depended on stain concentration, with the highest white-grey matter contrast (C1) occurring after 20 days at 1.25%, 22 days at 2.5%, and 15 days at 5.0% (Fig. 4a). At 5.0%, white-grey matter contrast declined after 15 days, consistent with progressive oversaturation of neural tissues. The maximum C1 contrast was similar across concentrations (∼0.17–0.20), indicating that stain concentrations primarily influence the time to achieve peak contrast rather than the maximum attainable value. Conversely, as staining continued and neural tissues became increasingly radiopaque, contrast between the brain and surrounding bone (C2) decreased and, in some specimens, reversed. These results demonstrate a clear trade-off between maximising internal neural contrast and maintaining brain-bone separation (Fig. 4b).

**Figure 4.**
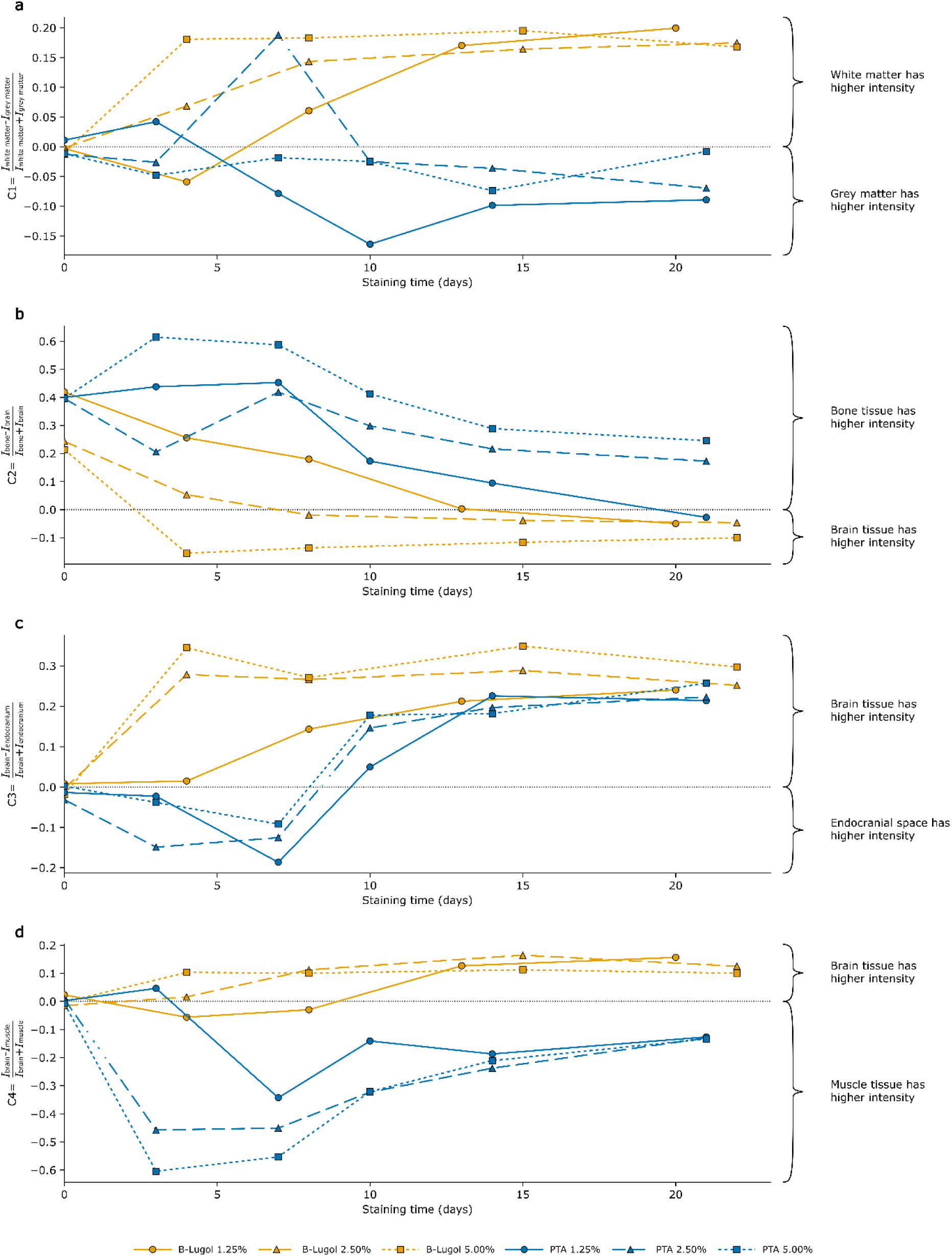
Time-dependent changes in tissue contrast across staining treatments. Signed Michelson contrast coefficients (C1, C2, C3 and C4) were calculated as *C_A_*_−*B*_ = (*I_A_* − *I_B_*)/(*I_A_* + *I_B_*). Positive values indicate that the first tissue has a higher greyscale intensity than the second; negative values indicate the reverse. Values near zero indicate similar intensities and limited differentiation, whereas larger absolute values indicate greater contrast, irrespective of polarity. (a) White–grey matter contrast (C1; white matter minus grey matter) is predominantly positive in buffered Lugol’s iodine-treated specimens. PTA-treated specimens exhibit values near zero or negative values, indicating similar tissue intensities or greater grey matter intensity, respectively. (b) Bone–brain contrast (C2; bone minus higher-intensity neural tissue) decreases during buffered Lugol’s iodine staining and becomes negative when neural tissue exceeds bone in greyscale intensity. PTA-treated specimens maintain positive values, indicating greater bone intensity. (c) Brain–endocranial space contrast (C3; lower-intensity neural tissue minus endocranial space) increases over time in both treatments, generally reaching higher values with buffered Lugol’s iodine. (d) Brain–muscle contrast (C4; higher-intensity neural tissue minus muscle) differs between treatments. Positive values indicate greater neural tissue intensity, whereas negative values indicate greater muscle intensity, reflecting the pronounced staining of musculature by PTA.

On the other hand, PTA did not exhibit a distinct optimal staining window for white-grey matter differentiation. white-grey matter contrast (C1) remained low across all time points and concentrations (Fig. 4a), indicating a limited ability to differentiate between these neural tissue types under the conditions tested. Despite this, contrast between bone and brain tissue remained positive and stable (Fig. 4b), while contrast between brain tissue and the endocranial space improved gradually over time (Fig. 4c). As staining progressed, muscular tissues became increasingly radiopaque, matching or even surpassing the intensity of the surrounding bone. This finding is consistent with the known affinity of PTA for protein-rich tissues (Metscher, 2009a; Pauwels *et al*., 2013; Quintarelli *et al*., 1971) and likely explains the strong contrast observed in surrounding musculature and tongue. Since PTA-treated specimens were imaged only until complete stain penetration had been achieved, we cannot exclude the possibility that longer incubation periods would further affect tissue contrast. Future studies should therefore investigate the taxon- and tissue-specific performance of PTA over extended staining durations, with particular emphasis on potential differences in staining affinity among muscle types, including skeletal and cardiac muscle (Doost and Arnolda, 2021).

Consistent with previous diceCT studies in reptiles (Callahan *et al*., 2021; Gignac *et al*., 2016; Hughes *et al*., 2016), our results show that buffered iodine staining produces white matter that is hyperintense (i.e. exhibits higher greyscale values and greater radiopacity), reflecting the preferential accumulation of iodine within lipid-rich myelinated fibre tracts (Gignac and Kley, 2018). Conversely, grey matter exhibits lower radiopacity, corresponding to regions dominated by neuronal cell bodies, dendrites, and relatively sparse myelinated fibres (Balcaen *et al*., 2025). In the brain of *A. muricatus*, examples of conspicuously hyperintense structures following iodine staining include the optic chiasm and optic tracts, the fibre-rich tectal layers, and cerebellar white matter (Figs. 5 and 6). Structures of intermediate intensity include the telencephalic dorsal ventricular ridge, cortical layers I and III, and the cerebellar molecular layer, whereas less radiopaque structures include cortical layer II and the cerebellar granular layer.

**Figure 5.**
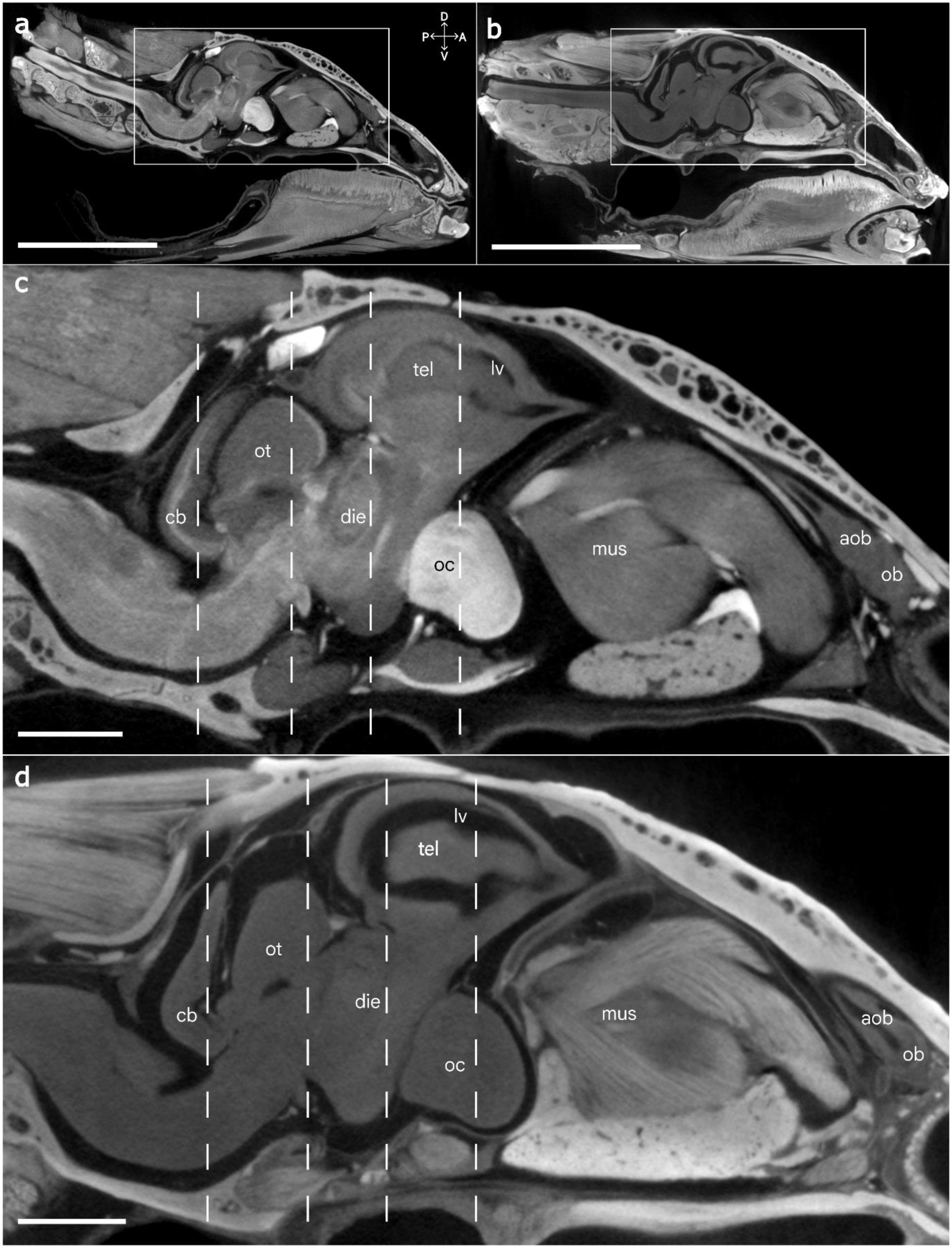
µCT virtual sagittal sections of a 1.25% buffered Lugol’s iodine-stained specimen (a, c), and a PTA-stained specimen (b, d) of *A. muricatus*. Iodine and PTA selectively bind to different molecules naturally present in soft tissues. These affinities are expressed as differential X-ray attenuation by specific tissues, generating distinct contrast patterns. Panels c and d correspond to the rectangles in a and b, respectively. Vertical dashed lines indicate the approximate locations of the coronal virtual sections shown in Fig. 6. aob, accessory olfactory bulb; cb, cerebellum; die, diencephalon; lv, lateral ventricle; mus; extraocular muscle; ob, olfactory bulb; oc, optic chiasm; ot, optic tectum; tel, telencephalon. Scale bar a, b = 1 cm; c,d = 2 mm.

**Figure 6.**
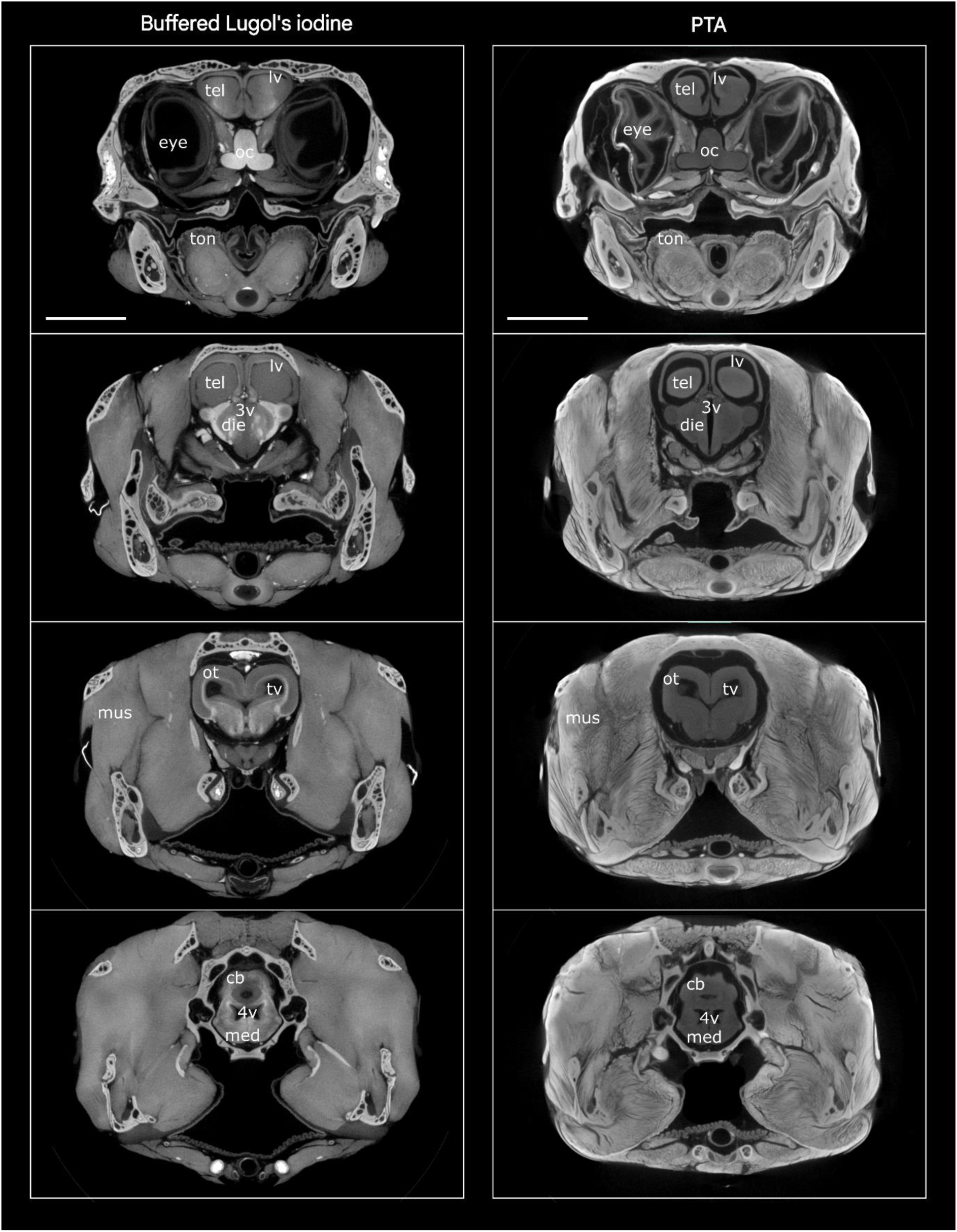
µCT series of virtual coronal sections of buffered Lugol’s iodine-stained and PTA-stained specimens of *A. muricatus*. Buffered Lugol’s iodine provides greater contrast between white matter and grey matter, whereas PTA enhances contrast in musculoskeletal tissue. Apparent brain shrinkage and ventricular expansion are evident in the PTA-stained specimen. 3v, third ventricle; 4v, fourth ventricle; cb, cerebellum; die, diencephalon; lv, lateral ventricle; med, medulla oblongata; mus; extraocular muscle; oc, optic chiasm; ot, optic tectum; tel, telencephalon; tv, tectal ventricle. Scale bars = 1 cm.

This level of tissue differentiation expands the utility of diceCT by enabling the identification of regional and layer-specific variation in neural organisation. It also demonstrates the potential for layer-specific or cell density-informed segmentation of selected brain regions in high-resolution scans, while retaining their three-dimensional anatomical relationships (Fig. 7). For example, variation in the relative volumes of individual cortical layers, cerebellar layers, or sensory-processing nuclei, such as the nucleus sphericus, could be examined in relation to sensory ecology, locomotor specialisations, or species-specific behavioural adaptations.

**Figure 7.**
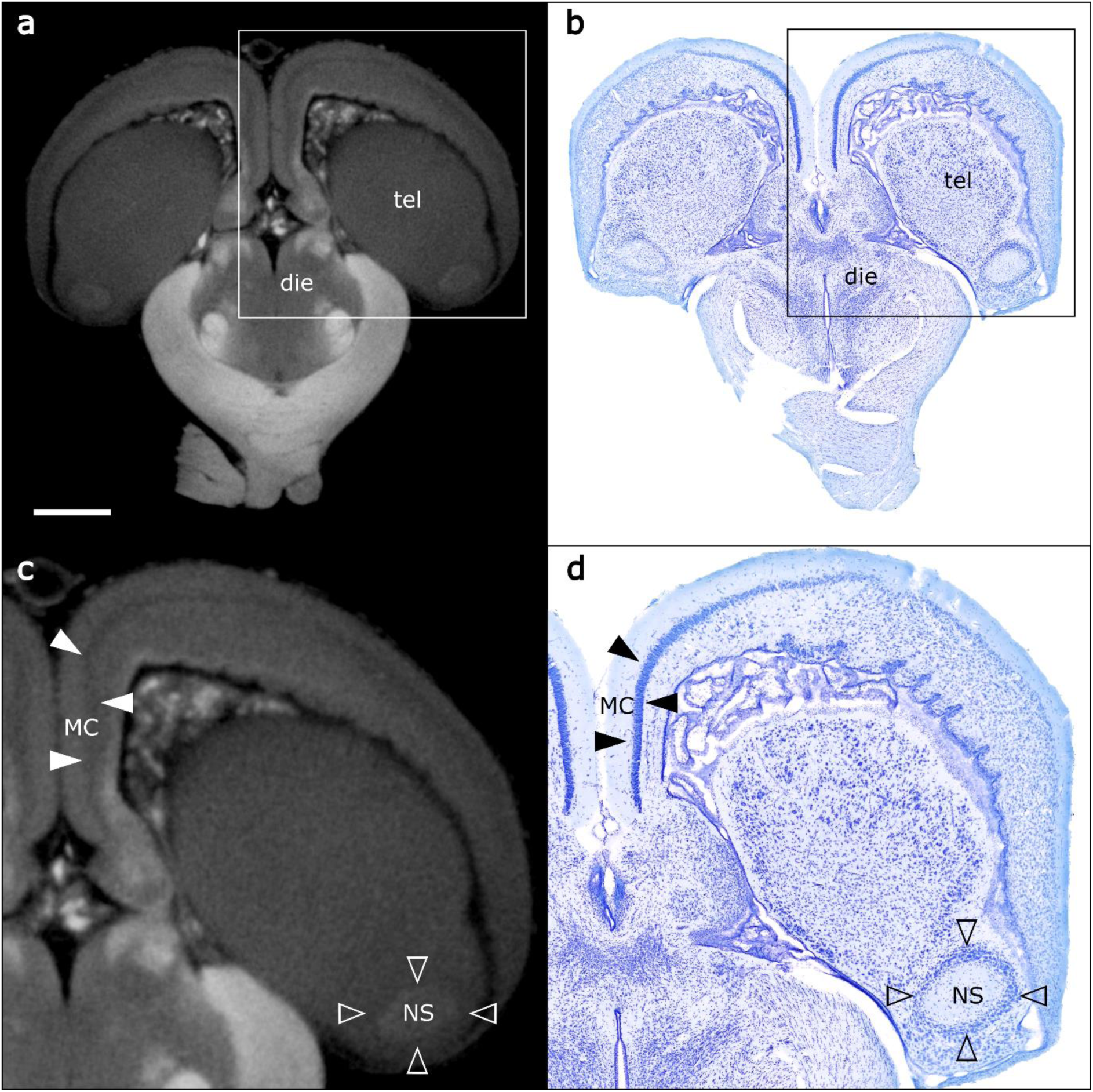
Comparison of corresponding coronal sections of the brain of *A. muricatus* visualised using diceCT and histology. Virtual µCT sections of the brain of a specimen stained with buffered Lugol’s iodine and scanned at a voxel size of 10 µm are shown in (a, c); cresyl violet-stained histological sections are shown in (b, d). The enlarged views (c, d) demonstrate that diceCT can resolve cortical layers (e.g., layer II of the medial cortex, indicated by solid arrowheads) and sensory nuclei (e.g., the nucleus sphericus, indicated by open arrowheads), while largely preserving their native spatial relationships. die, diencephalon; MC, medial cortex; NS, nucleus sphericus; tel, telencephalon. Scale bar = 1 mm.

### Brain occupancy, tissue shrinkage and brain-endocast correspondence

Quantitative assessment of brain occupancy within the endocranial cavity suggests considerably less shrinkage in specimens stained with buffered Lugol’s iodine than in those stained with PTA (Fig. 8 and Supplementary File S3), regardless of staining agent concentration. All descriptive values are reported as mean ± SD. Buffered Lugol’s iodine specimens exhibited greater brain occupancy, with a mean brain-to-endocast ratio (BER) of 0.708 ± 0.020 (n = 4) compared with 0.614 ± 0.032 (n = 5) for PTA-treated specimens. This pattern remained after exclusion of the ventricular system, with mean brain tissue-to-endocast ratio (BTER) values of 0.695 ± 0.017 and 0.558 ± 0.024 for buffered Lugol’s iodine and PTA, respectively. Consistent with these results, the ventricular system occupied a substantially smaller proportion of total brain volume in iodine-stained specimens (1.84% ± 0.46%) than in PTA-stained specimens (9.19% ± 0.91%), suggesting greater relative ventricular enlargement associated with shrinkage of the surrounding periventricular brain tissue following PTA treatment.

**Figure 8.**
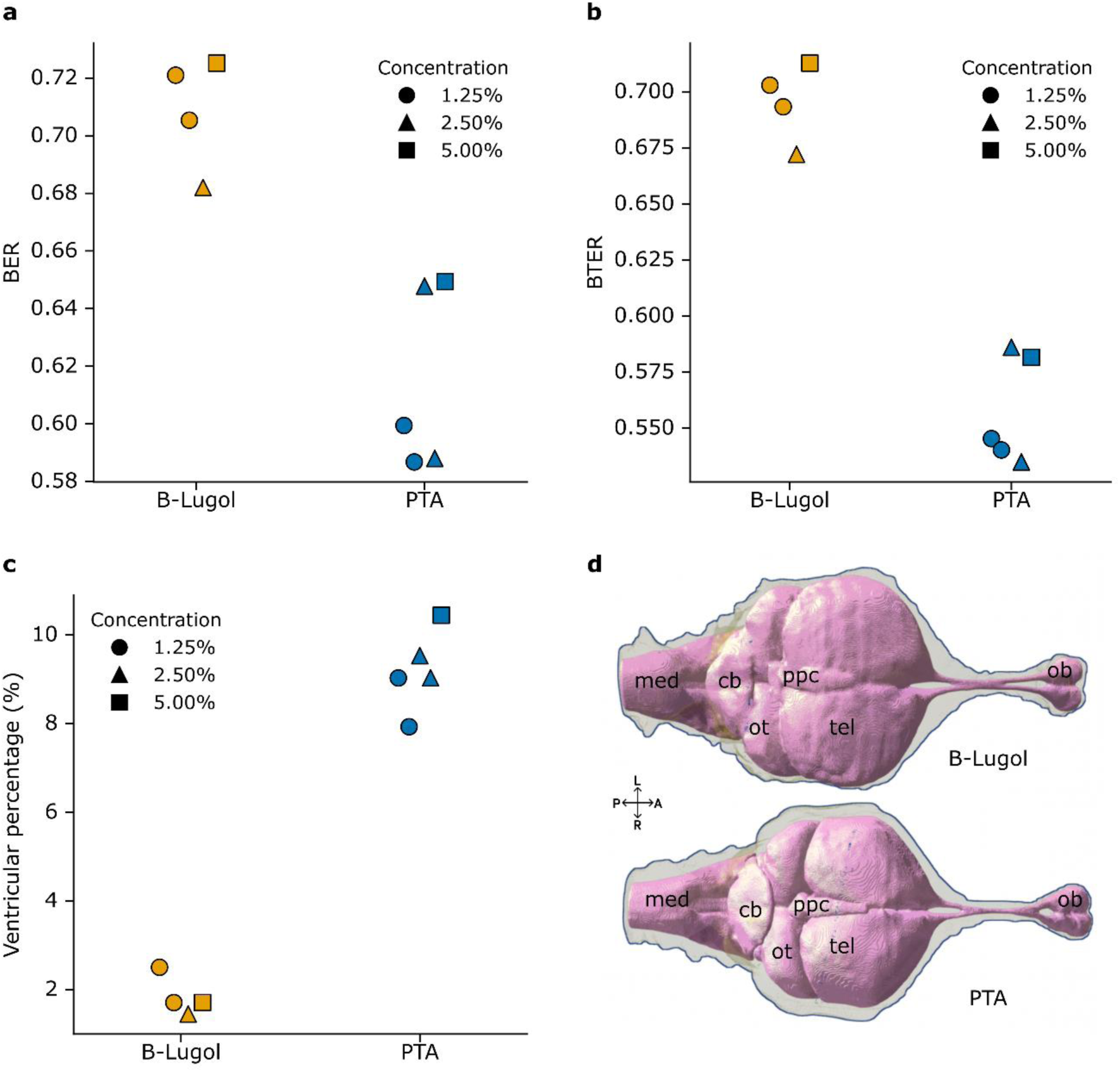
Brain-to-endocast ratio (BER) (a), brain tissue-to-endocast ratio (BTER) (b), and brain ventricular percentage (c) in buffered Lugol’s iodine-stained (orange symbols) and PTA-stained specimens (blue symbols). PTA-induced brain tissue shrinkage results in reduced occupancy of the endocranial cavity. Symbol shapes represent staining concentrations. (d) Virtual segmentations of the brain (pink) and endocast (grey) of a buffered Lugol’s-stained (top) and a PTA-stained specimen (bottom) in dorsal view. Brain shrinkage is visible as a greater separation between the external surface of the brain and the internal surface of the endocranial cavity, especially noticeable in the telencephalon and optic tectum. cb, cerebellum; med, medulla oblongata; ob, olfactory bulb; ot, optic tectum; ppc, parapineal complex; tel, telencephalon.

Differences between contrast agents persisted after separate adjustment for body mass, SVL, head width, or endocast volume, with no evidence of associations between these covariates and the outcomes (Supplementary File S3). No consistent relationship between stain concentration and BER or BTER was observed within either treatment, although the small sample sizes limited assessment of concentration effects. Specimens were therefore pooled by contrast agent for the primary comparison.

A reasonable question arising from these differences in brain-to-endocast occupancy concerns the spatial pattern of tissue shrinkage. How does brain shrinkage affect the morphological correspondence between the brain and endocast? Our cloud-to-cloud comparisons between the brain and endocast provide a visual representation of their surface morphological correspondence (Fig. 9). Regardless of the staining treatment, three-dimensional heat maps of surface distances revealed regions of particularly close brain-endocast fit, including the olfactory bulbs, parts of the cerebral hemispheres, the pineal organ, and the hypophysis. Conversely, certain anatomical regions exhibited poorer brain-endocast correspondence, most notably the cerebellum and the rostral telencephalon (Figs. 9-11). Overall, buffered Lugol’s-stained specimens exhibited greater and more spatially consistent correspondence between the brain and endocast, reflected by a larger proportion of regions with minimal surface separation, particularly within the forebrain (Fig. 10). In contrast, PTA-stained specimens showed a greater proportion of intermediate to large surface distances with heterogeneous shrinkage across the whole brain with greater regional variation in the dorsal and ventrolateral regions of the forebrain (Fig. 11). These results have implications for studies that use contrast-enhanced imaging for determining brain shape and volume (e.g., Camilieri-Asch *et al*., 2020b; Macrì *et al*., 2019) and the occupancy of the brain within the cranium (e.g., Allemand *et al*., 2023; Clement *et al*., 2015; Figueroa and Pierce, 2026; Watanabe *et al*., 2019).

**Figure 9.**
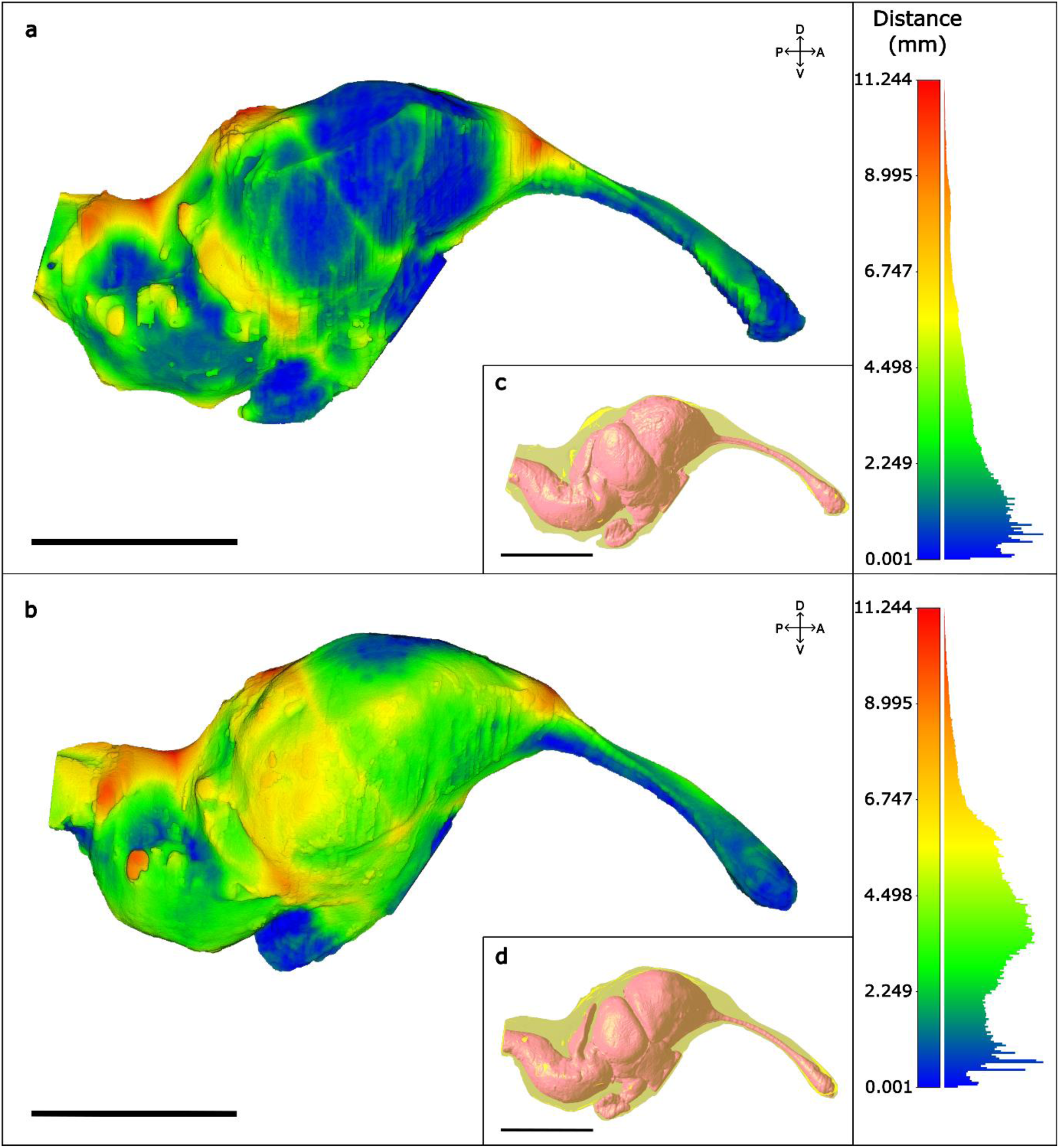
Cloud to cloud comparison between the brain and endocast of a buffered Lugol’s iodine-stained specimen (a) and a PTA-stained (b) specimen, together with histograms showing the corresponding surface-distance distributions. Low distance values (cool colours) indicate areas where the brain closely fits to the endocranial cavity, whereas higher distance values (warm colours) represent increased separation between the brain and endocast. The predominance of low distance values in (a) suggests closer shape correspondence between the brain and endocast, whereas the higher frequency of intermediate values in (b) implies greater regional variation in brain-endocast correspondence, consistent with brain tissue shrinkage. (c) and (d) illustrate the position of brain within the braincase for the specimens shown in (a) and (b), respectively. Scale bars = 0.5 cm.

**Figure 10.**
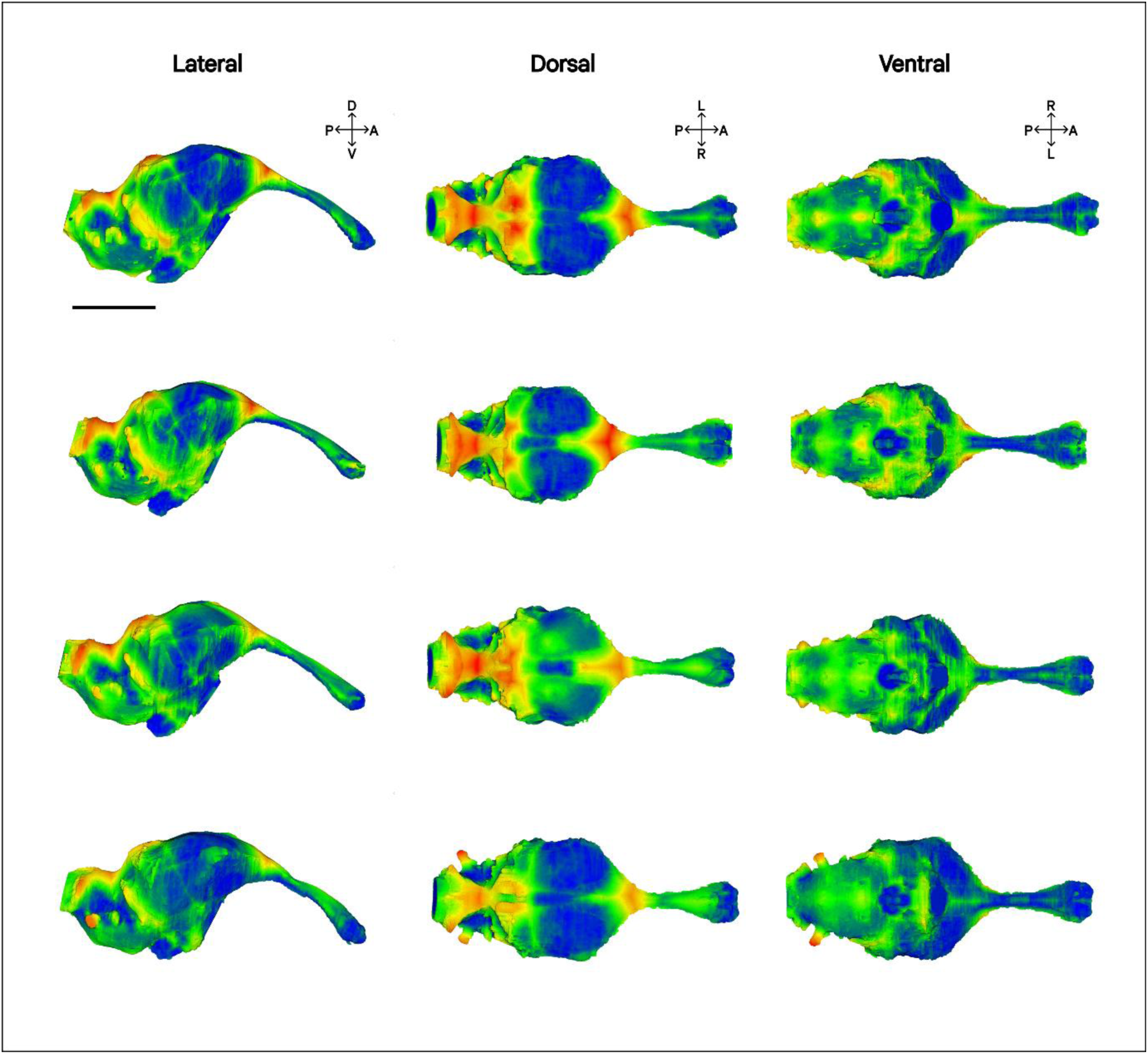
Cloud to cloud comparison between the brain and endocast of buffered Lugol’s iodine-stained specimens of *A. muricatus*. Each row corresponds to a different specimen. Cool colours represent areas where the brain closely fits to the endocranial cavity, whereas warm colours represent greater separation between the brain and endocast. The distance maps were dominated by low absolute distance values, indicating close brain-endocast correspondence, particularly in the forebrain. Scale bar = 0.5 cm.

**Figure 11.**
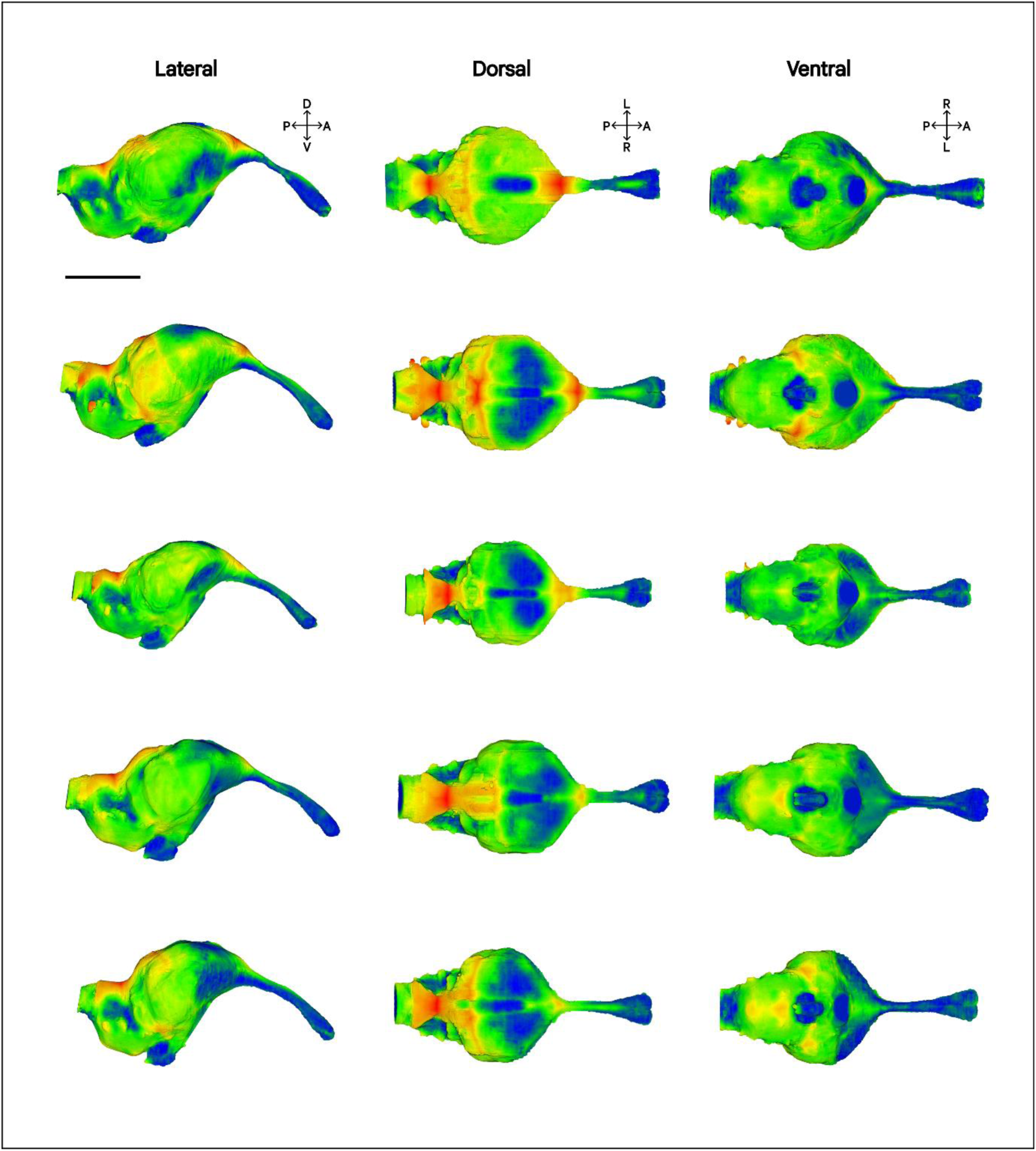
Cloud to cloud comparison between the brain and endocast of PTA-stained specimens of *A. muricatus.* Each row corresponds to a different specimen. Cool colours indicate areas where the brain closely fit to the endocranial cavity, whereas warm colours reflect greater separation between the brain and endocast. PTA-stained specimens showed broader distributions with increased frequencies of intermediate surface distances, indicating reduced brain–endocast correspondence and greater regional variability. Scale bar = 0.5 cm.

Tissue shrinkage remains one of the principal limitations of contrast-enhanced µCT using both iodine- and PTA-based stains (Buytaert *et al*., 2014; Dawood *et al*., 2021; Procházková *et al*., 2026; Vickerton et al., 2013). We adopted two approaches to quantify shrinkage by defining the brain-to-endocast ratio (BER) and brain tissue-to-endocast ratio (BTER), which respond to different aspects of tissue shrinkage. BER evaluates changes in the occupancy of the brain as an organ within the endocranial cavity following contrast-enhancement staining of soft tissues, making it particularly relevant to geometric morphometric studies. Our results show a difference of 9.4 percentage points in brain occupancy between treatments, corresponding to a 13.3% lower occupancy in PTA-treated specimens than in buffered Lugol’s iodine-stained specimens. In contrast, BTER assesses how the neural tissue constituting the brain responds to the different staining procedures, providing a comparative measure of how contrast-enhancing agents impact soft-tissue preservation. BTER was 13.7 percentage points higher in buffered Lugol’s specimens than in PTA specimens, which is consistent with approximately 20% greater apparent brain tissue shrinkage relative to buffered Lugol’s specimens. Nevertheless, since the true *in vivo* brain-to-endocast relationship is not known for this species, these values should be interpreted comparatively rather than as absolute measures of shrinkage. It is also worth noting that our specimens were fixed in buffered fixative and stored in buffer prior to treatments and therefore were likely to have experienced less shrinkage compared to ethanol-preserved museum specimens that are typically used in comparative diceCT studies (Hedrick *et al*., 2018).

Most studies addressing soft-tissue shrinkage artefacts associated with iodine or PTA staining predate Dawood *et al*. (2021), who demonstrated that buffered Lugol’s solution can substantially reduce soft-tissue shrinkage (from 31.9% to 5.9% in mouse livers) without compromising staining efficacy relative to unbuffered formulations. Those earlier studies reported that PTA induces less shrinkage than unbuffered iodine-based stains, suggesting PTA as a preferable option when preservation of original tissue volume is a critical consideration (Buytaert *et al*., 2014; Pauwels *et al*., 2013). In contrast to these findings, our results indicate that iodine, when applied as buffered Lugol’s iodine, produces less apparent brain tissue shrinkage than aqueous PTA by approximately 20% under the conditions tested. Furthermore, to our knowledge, this study is the first to quantitatively assess both total brain (including ventricular system) and brain tissue shrinkage following contrast-enhanced staining, demonstrating that buffered Lugol’s iodine improves white matter–grey matter contrast while reducing brain shrinkage relative to PTA under the conditions tested.

Buffered Lugol’s iodine is a practical alternative to unbuffered Lugol’s iodine because it provides pH stability, as acidification, rather than osmolarity, has been proposed as the primary factor contributing to tissue shrinkage during iodine treatment (Dawood *et al*., 2021). The greater shrinkage observed in PTA-stained specimens in this study, relative to those stained with buffered Lugol’s iodine, may be related to the typically low pH reported in aqueous PTA solutions. The unadjusted PTA solutions used in the primary comparison had pH values below 2.0, which may contribute, at least in part, to the reduction in brain tissue volume.

Since relatively little information is available on strategies to reduce shrinkage artefacts during PTA staining, we conducted an exploratory assessment of PTA solutions adjusted to different pH levels, including mildly acidic (pH 4.0) and neutral (pH 7.2) conditions. Qualitative observations suggest that increasing solution pH reduced tissue shrinkage, as indicated by a smaller separation between the dorsal surface of the brain and the endocranial roof, together with less apparent expansion of the ventricular system compared with the PTA-treated specimens used in the buffered Lugol’s iodine comparison. However, these apparent improvements in tissue preservation were accompanied by lower overall soft-tissue contrast, poorer image quality, and substantially longer staining times, particularly for the neutral pH solution, which was evaluated after six months of staining (Supplementary Fig. S2), suggesting a potential trade-off between tissue integrity and staining efficiency. This pattern is consistent with the staining mechanism of PTA, which binds preferentially to positively charged protein groups that become increasingly protonated below their isoelectric point, resulting in stronger tissue interactions under acidic conditions (Quintarelli *et al*., 1971). In this exploratory assessment, aqueous PTA at pH 4 appeared to reduce shrinkage while retaining sufficient contrast to distinguish the overall brain outline. Further systematic studies are required to determine whether other pH ranges yield similar or improved outcomes and whether buffer incorporation can effectively mitigate PTA-induced tissue shrinkage while maintaining sufficient contrast enhancement for µCT imaging.

## Conclusions

To our knowledge, this study provides the first quantitative comparison of contrast quality and brain tissue shrinkage between buffered Lugol’s iodine- and aqueous PTA staining for brain CT imaging in vertebrates. Under the conditions tested, we show that buffered Lugol’s iodine was the more effective of the two staining protocols for detailed brain µCT characterisation in *A. muricatus*, with potential applications in other vertebrates. We recommend using buffered Lugol’s iodine when the primary objective is to maximise internal neural contrast for fine-scale anatomical investigations in vertebrates or to preserve brain morphology for volumetric and shape-based analyses, whereas PTA offers advantages for general soft-tissue discrimination or musculoskeletal studies in vertebrates. Finally, to ensure more accurate comparative neuroanatomical studies using µCT-generated datasets, we recommend specifying the fixation and preservation method, contrast agent and concentration, and staining protocol employed. This will help account for volumetric and/or morphological artefacts, such as those reported here, when comparing specimens prepared using different contrast-enhancement techniques and preserved using different storage methods, particularly when comparing freshly fixed specimens with ethanol-preserved museum specimens.

## Author contributions

F.N., J.M.C-R. and S.P.C. conceived of the study, S.W. and D.S.B.D. provided the specimens; F.N. conducted experiments and analysed the data with assistance from C.C.K.; J.R.B. performed the microCT scanning and reconstructions with assistance from F.N.; F.N. wrote the manuscript with input from all co-authors.

## Acknowledgements

We thank Kushini Kularatne for specimen collection; Dr Lucille Chapuis for providing software and laboratory support; Dr Jacquie Orian for kindly providing histological materials; Dr Chad Johnson from La Trobe University’s Bioimaging Platform for assistance with digitising the histological material; the Melbourne Trace Analysis for Chemical, Earth and Environmental Sciences (TrACEES) Platform for access to the micro-CT; Dr Remi Allemand and Dr Daniel Hoops for advice on brain and endocast segmentation; and Dr Amanda Ibsen for advice on statistical analyses. ChatGPT (OpenAI) was used for grammar, spelling and readability checks. The authors reviewed all output and take full responsibility for the final content.

## Ethics statement

All procedures adhered to La Trobe University and University of Canberra animal ethics guidelines (Project 270) and complied with national welfare guidelines.

## Data accessibility

The data used in this study are provided as electronic supplementary material. Micro-CT data will be uploaded to MorphoSource upon publication (Project ID: 000872072).

