## Supplementary File S1 for "Optimising neural contrast and minimising tissue shrinkage in dragon lizard brains: buffered Lugol’s iodine versus phosphotungstic acid for contrast-enhanced µCT"

Supplementary Table 1. Contrast enhancing and scanning parameters tested per specimen

| Specimen ID | Scan  number | Staining  agent | Concentration | Solvent | Solution pH | Staining  Time  (days) | Scan  Time  (min) | Voltage  (kV) | Current  (mA) | Resolution  (µm) | Timing  (msec) | Images | Filter |
| --- | --- | --- | --- | --- | --- | --- | --- | --- | --- | --- | --- | --- | --- |
| AMLatrobe1 | 1 | None | - | - | - | 0 | 10 | 75 | 300 | 20 | 500 | 1199 | none |
| AMLatrobe1 | 2 | Lugol's iodine | 1.25% | Sorensen's buffer | 7.2 | 4 | 10 | 75 | 300 | 20 | 500 | 1199 | none |
| AMLatrobe1 | 3 | Lugol's iodine | 1.25% | Sorensen's buffer | 7.2 | 8 | 10 | 75 | 300 | 20 | 500 | 1199 | none |
| AMLatrobe1 | 4 | Lugol's iodine | 1.25% | Sorensen's buffer | 7.2 | 13 | 10 | 75 | 300 | 20 | 500 | 1199 | none |
| AMLatrobe1 | 5 | Lugol's iodine | 1.25% | Sorensen's buffer | 7.2 | 20 | 10 | 75 | 300 | 20 | 500 | 1199 | none |
| AMLatrobe2 | 1 | None | - | - | - | 0 | 10 | 120 | 300 | 20 | 500 | 1199 | none |
| AMLatrobe2 | 2 | Lugol's iodine | 2.50% | Sorensen's buffer | 7.2 | 1 | 10 | 120 | 300 | 20 | 500 | 1199 | none |
| AMLatrobe2 | 3 | Lugol's iodine | 2.50% | Sorensen's buffer | 7.2 | 4 | 10 | 120 | 300 | 20 | 500 | 1199 | 0.1mm Cu |
| AMLatrobe2 | 4 | Lugol's iodine | 2.50% | Sorensen's buffer | 7.2 | 8 | 10 | 120 | 300 | 20 | 500 | 1199 | 0.1mm Cu |
| AMLatrobe2 | 5 | Lugol's iodine | 2.50% | Sorensen's buffer | 7.2 | 15 | 10 | 120 | 300 | 20 | 500 | 1199 | 0.1mm Cu |
| AMLatrobe2 | 6 | Lugol's iodine | 2.50% | Sorensen's buffer | 7.2 | 22 | 10 | 120 | 300 | 20 | 500 | 1199 | 0.1mm Cu |
| AMLatrobe3 | 1 | None | - | - | - | 0 | 10 | 120 | 300 | 20 | 500 | 1199 | none |
| AMLatrobe3 | 2 | Lugol's iodine | 5% | Sorensen's buffer | 7.2 | 1 | 10 | 120 | 300 | 20 | 500 | 1199 | none |
| AMLatrobe3 | 3 | Lugol's iodine | 5% | Sorensen's buffer | 7.2 | 4 | 10 | 120 | 300 | 20 | 500 | 1199 | 0.1mm Cu |
| AMLatrobe3 | 4 | Lugol's iodine | 5% | Sorensen's buffer | 7.2 | 8 | 10 | 120 | 300 | 20 | 500 | 1199 | 0.1mm Cu |
| AMLatrobe3 | 5 | Lugol's iodine | 5% | Sorensen's buffer | 7.2 | 15 | 10 | 120 | 300 | 20 | 500 | 1199 | 0.1mm Cu |
| AMLatrobe3 | 6 | Lugol's iodine | 5% | Sorensen's buffer | 7.2 | 22 | 10 | 120 | 300 | 20 | 500 | 1199 | 0.1mm Cu |
| AMLatrobe4 | 1 | None | - | - | - | 0 | 10 | 120 | 300 | 20 | 500 | 1199 | none |
| AMLatrobe4 | 2 | PTA | 1.25% | Water | 4 | 1 | 10 | 120 | 300 | 20 | 500 | 1199 | none |
| AMLatrobe4 | 3 | PTA | 1.25% | Water | 4 | 4 | 10 | 120 | 300 | 20 | 500 | 1199 | 0.1mm Cu |
| AMLatrobe4 | 4 | PTA | 1.25% | Water | 4 | 8 | 10 | 120 | 300 | 20 | 500 | 1199 | 0.1mm Cu |
| AMLatrobe4 | 5 | PTA | 1.25% | Water | 4 | 15 | 10 | 120 | 300 | 20 | 500 | 1199 | 0.1mm Cu |
| AMLatrobe4 | 6 | PTA | 1.25% | Water | 4 | 22 | 10 | 120 | 300 | 20 | 500 | 1199 | 0.1mm Cu |
| AMLatrobe5 | 1 | None | - | - | - | 0 | 10 | 120 | 300 | 20 | 500 | 1199 | none |
| AMLatrobe5 | 2 | PTA | 2.50% | Water | 4 | 1 | 10 | 120 | 300 | 20 | 500 | 1199 | none |
| AMLatrobe5 | 3 | PTA | 2.50% | Water | 4 | 4 | 10 | 120 | 300 | 20 | 500 | 1199 | 0.1mm Cu |
| AMLatrobe5 | 4 | PTA | 2.50% | Water | 4 | 8 | 10 | 120 | 300 | 20 | 500 | 1199 | 0.1mm Cu |
| AMLatrobe5 | 5 | PTA | 2.50% | Water | 4 | 15 | 10 | 120 | 300 | 20 | 500 | 1199 | 0.1mm Cu |
| AMLatrobe5 | 6 | PTA | 2.50% | Water | 4 | 22 | 10 | 120 | 300 | 20 | 500 | 1199 | 0.1mm Cu |
| AMLatrobe7 | 1 | None | - | - | - | 0 | 10 | 75 | 300 | 20 | 500 | 1199 | none |
| AMLatrobe7 | 2 | Lugol's iodine | 1.25% | Sorensen's buffer | 7.2 | 2 | 10 | 75 | 300 | 20 | 500 | 1199 | none |
| AMLatrobe7 | 3 | Lugol's iodine | 1.25% | Sorensen's buffer | 7.2 | 6 | 10 | 75 | 300 | 20 | 500 | 1199 | none |
| AMLatrobe7 | 4 | Lugol's iodine | 1.25% | Sorensen's buffer | 7.2 | 11 | 10 | 75 | 300 | 20 | 500 | 1199 | none |
| AMLatrobe7 | 5 | Lugol's iodine | 1.25% | Sorensen's buffer | 7.2 | 18 | 10 | 75 | 300 | 20 | 500 | 1199 | none |
| AMLatrobe7 | 6 | Lugol's iodine | 1.25% | Sorensen's buffer | 7.2 | 45 | 10 | 75 | 300 | 20 | 500 | 1199 | none |
| AMLatrobe8 | 1 | None | - | - | - | 0 | 10 | 120 | 300 | 20 | 500 | 1199 | 0.1mm Cu |
| AMLatrobe8 | 2 | PTA | 2.50% | Water | 2 | 1 | 10 | 120 | 300 | 20 | 500 | 1199 | 0.1mm Cu |
| AMLatrobe8 | 3 | PTA | 2.50% | Water | 2 | 5 | 10 | 120 | 300 | 20 | 500 | 1199 | 0.1mm Cu |
| AMLatrobe8 | 4 | PTA | 2.50% | Water | 2 | 8 | 10 | 120 | 300 | 20 | 500 | 1199 | 0.1mm Cu |
| AMLatrobe8 | 5 | PTA | 2.50% | Water | 2 | 12 | 10 | 120 | 300 | 20 | 500 | 1199 | 0.1mm Cu |
| AMLatrobe8 | 6 | PTA | 2.50% | Water | 2 | 20 | 10 | 120 | 300 | 20 | 500 | 1199 | 0.1mm Cu |
| AMLatrobe12 | 1 | None | - | - | - | 0 | 10 | 120 | 300 |  | 500 | 1199 | none |
| AMLatrobe12 | 2 | PTA | 1.25% | Water | 7.2 | 6 | 10 | 120 | 300 | 20 | 500 | 1199 | none |
| AMLatrobe12 | 3 | PTA | 1.25% | Water | 7.2 | 14 | 10 | 120 | 300 | 20 | 500 | 1199 | none |
| AMLatrobe12 | 4 | PTA | 1.25% | Water | 7.2 | 153 | 10 | 120 | 300 | 20 | 500 | 1199 | none |
| AMLatrobe14 | 1 | None | - | - | - | 0 | 10 | 120 | 300 | 20 | 500 | 1199 | none |
| AMLatrobe14 | 2 | PTA | 1.25% | Water | 7.2 | 6 | 10 | 120 | 300 | 20 | 500 | 1199 | none |
| AMLatrobe14 | 3 | PTA | 1.25% | Water | 7.2 | 14 | 10 | 120 | 300 | 20 | 500 | 1199 | none |
| 81820 | 1 | None | - | - | - | 0 | 10 | 120 | 300 | 20 | 500 | 1199 | 0.1mm Cu |
| 81820 | 2 | PTA | 5% | Water | 2 | 1 | 10 | 120 | 300 | 20 | 500 | 1199 | 0.1mm Cu |
| 81820 | 3 | PTA | 5% | Water | 2 | 5 | 10 | 120 | 300 | 20 | 500 | 1199 | 0.1mm Cu |
| 81820 | 4 | PTA | 5% | Water | 2 | 8 | 10 | 120 | 300 | 20 | 500 | 1199 | 0.1mm Cu |
| 81820 | 5 | PTA | 5% | Water | 2 | 12 | 10 | 120 | 300 | 20 | 500 | 1199 | 0.1mm Cu |
| 81820 | 6 | PTA | 5% | Water | 2 | 20 | 10 | 120 | 300 | 20 | 500 | 1199 | 0.1mm Cu |
| 81821 | 1 | None | - | - | - | 0 | 10 | 75 | 300 | 20 | 500 | 1199 | none |
| 81821 | 2 | PTA | 1.25% | Water | 2 | 22 | 10 | 120 | 300 | 20 | 500 | 1199 | none |
| 81928 | 1 | None | - | - | - | 0 | 10 | 75 | 300 | 20 | 500 | 1199 | none |
| 81928 | 2 | PTA | 2.50% | Water | 2 | 22 | 10 | 120 | 300 | 20 | 500 | 1199 | none |
| 81928 | 3 | PTA | 2.50% | Water | 2 | 34 | 10 | 120 | 300 | 20 | 500 | 1199 | none |
| 9512 | 1 | None | - | - | - | 0 | 10 | 120 | 300 | 20 | 500 | 1199 | 0.1mm Cu |
| 9512 | 2 | PTA | 1.25% | Water | 2 | 1 | 10 | 120 | 300 | 20 | 500 | 1199 | 0.1mm Cu |
| 9512 | 3 | PTA | 1.25% | Water | 2 | 5 | 10 | 120 | 300 | 20 | 500 | 1199 | 0.1mm Cu |
| 9512 | 4 | PTA | 1.25% | Water | 2 | 8 | 10 | 120 | 300 | 20 | 500 | 1199 | 0.1mm Cu |
| 9512 | 5 | PTA | 1.25% | Water | 2 | 12 | 10 | 120 | 300 | 20 | 500 | 1199 | 0.1mm Cu |
| 9512 | 6 | PTA | 1.25% | Water | 2 | 20 | 10 | 120 | 300 | 20 | 500 | 1199 | 0.1mm Cu |
