## Supplementary Figure S1 for "Optimising neural contrast and minimising tissue shrinkage in dragon lizard brains: buffered Lugol’s iodine versus phosphotungstic acid for contrast-enhanced µCT"

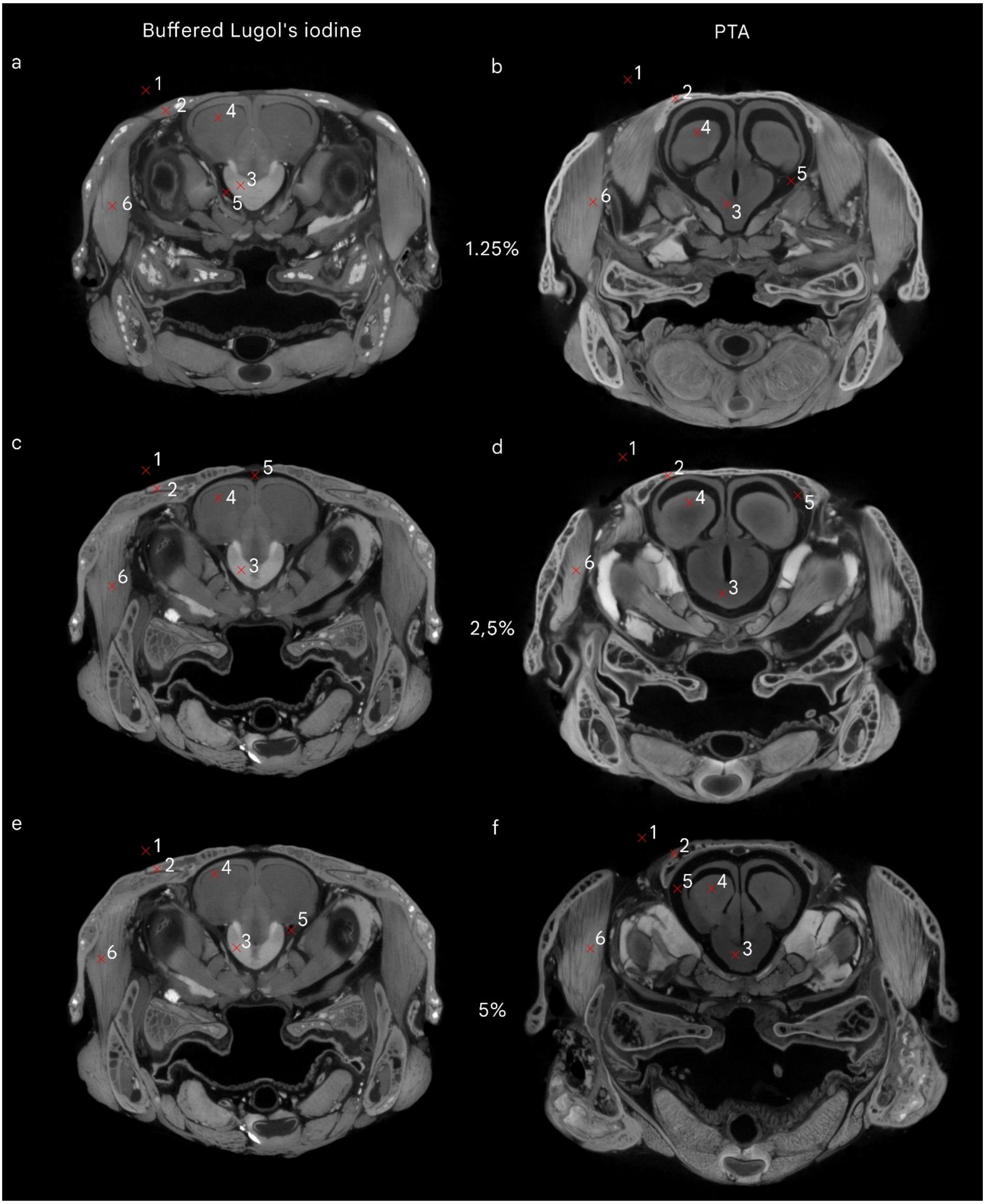


*Supplementary Figure S1. Representative virtual sections showing the anatomical locations of the six probe points placed in each buffered Lugol's-stained specimen (a, c, e) and PTA-stained specimen (b, d, f). The reconstructed TIFF stacks for each specimen were manually aligned, and greyscale values were extracted from all aligned datasets, ensuring that identical three-dimensional locations were sampled throughout the staining time series. Each probe point sampled a different tissue type or region of interest within the same section: background (1), bone (2), white matter (3), grey matter (4), endocranial space (5), and muscle (6).*
