## Supplementary Figure S2 for "Optimising neural contrast and minimising tissue shrinkage in dragon lizard brains: buffered Lugol’s iodine versus phosphotungstic acid for contrast-enhanced µCT"

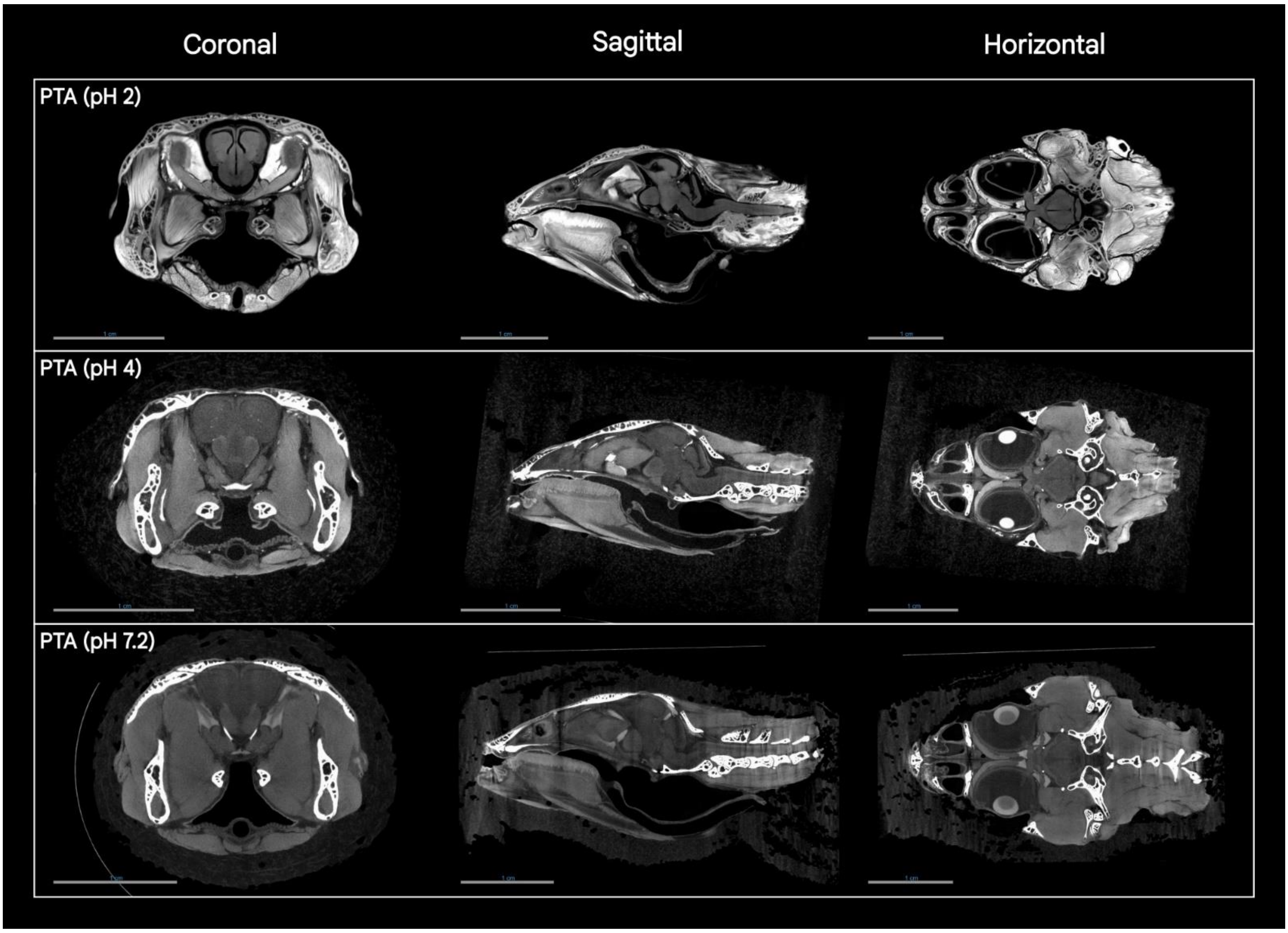


*Supplementary Figure S2. Effects of solution pH on contrast, image quality, and brain shrinkage in specimens of A. muricatus stained with a 1.25% aqueous PTA solution adjusted to three pH values: acidic (pH 2, 3 weeks post-staining; top row), mildly acidic (pH 4, 3 weeks post staining; middle row) and neutral (pH 7.2, 6 months post staining; bottom row). Increasing the solution pH reduced tissue shrinkage, as indicated by the smaller separation between the external surface of the brain and the internal surface of the endocranial cavity at pH 4 and 7.2, together with less apparent expansion of the ventricular system compared with the specimen treated with PTA at pH 2. The apparent improvement in tissue preservation was accompanied by lower overall soft tissue contrast, poorer image quality, and substantially longer staining times, particularly with the neutral solution. These observations suggest a potential trade-off between tissue integrity and staining efficiency. Scale bars = 1 cm.*
